# NFAT Transcription Factors are Essential and Redundant Actors for Leukemia Initiating Potential in T-cell Acute Lymphoblastic Leukemia

**DOI:** 10.1101/2020.10.30.362376

**Authors:** Claire Catherinet, Diana Passaro, Stéphanie Gachet, Hind Medyouf, Anne Reynaud, Charlène Lasgi, Jacques Ghysdael, Christine Tran Quang

**Author notes:** These authors contributed equally to this work (Claire Catherinet, Diana Passaro, Stéphanie Gachet).

## Abstract

T-cell acute lymphoblastic leukemia (T-ALL) is an aggressive malignancy with few available targeted therapies. We previously reported that the phosphatase calcineurin (Cn) is required for LIC (leukemia Initiating Capacity) potential of T-ALL pointing to Cn as an interesting therapeutic target. Calcineurin inhibitors have however unwanted side effect. NFAT transcription factors play crucial roles downstream of calcineurin during thymocyte development, T cell differentiation, activation and anergy. Here we elucidate NFAT functional relevance in T-ALL. Using murine T-ALL models in which *Nfat* genes can be inactivated either singly or in combination, we show that NFATs are required for T-ALL LIC potential and essential to survival, proliferation and migration of T-ALL cells. We also demonstrate that *Nfat* genes are functionally redundant in T-ALL and identified a node of genes commonly deregulated upon Cn or NFAT inactivation, which may serve as future candidate targets for T-ALL.

## INTRODUCTION

T-cell acute lymphoblastic leukemia (T-ALL) is an aggressive malignancy of T-cell progenitors that represents about 15% of pediatric and 25% of adult acute lymphoblastic leukemia cases. Genome-wide transcriptional profiling analysis enabled to classify T-ALL into different molecular subgroups characterized by abnormal expression of several transcription factors including TAL1/2, LMO1/2, TLX1/3, NKX2.1/2.2, HOXA, as the result of genetic rearrangements or other modes of deregulation (for review^1^). Another T-ALL subgroup encompasses cases characterized by a transcriptional signature resembling that of early T cell progenitors (ETP subgroup). Across these subgroups, a number of additional, recurrent alterations are found in tumor suppressor genes/loci, including *CDKN2A* or *PTEN* and in oncogenes, most notably *NOTCH1*, which harbors functionally relevant activating mutations in the majority of T-ALL cases^2, 3^

Besides genetic and epigenetic oncogenic cues, T-ALL development also depends upon specific micro-environmental signals (for review^4^). Recent evidence has shown that bone marrow (BM)-stroma produced CXCL12 acting through its CXCR4 receptor on T-ALL cells is essential to leukemia development and initiating potential^5, 6^. We notably found that cell-surface expression of CXCR4 in T-ALL depends upon the activation of calcineurin (PPP3, named Cn thereafter)^6^, a calcium-dependent phosphatase that we previously showed to be critical to T-ALL cell survival, proliferation, migratory activity and leukemia initiating potential^7, 8^. However, these studies also showed that restoring normal CXCR4 cell surface expression in Cn-deficient T-ALL failed to correct their impaired leukemia initiating potential^6^, indicating the existence of other Cn effectors critical to T-ALL biology.

NFAT transcription factors are important effectors of calcium/calcineurin signaling in normal T cell development and in many aspects of mature T cell functions (for review,^9^). NFAT factors are composed of a DNA binding domain structurally related to that of the REL/NFkB family protein, a regulatory domain in which 12-14 serine residues located in specific regions (SRR1; SP1-SP3) are targeted for phosphorylation and N- and C-terminal activation domains^9^. In unstimulated cells, NFAT proteins are hyperphosphorylated in their SRR/SP motifs by the cooperative action of several protein kinases, including glycogen synthase kinase 3 (GSK3), casein kinase 1 (CK1) and dual specificity tyrosine phosphorylation-regulated kinases (DYRK)^9^ and are sequestered as a supramolecular cytoplasmic complex^10^. Stimuli increasing calcium concentration and resulting in calcineurin activation lead to dephosphorylation of NFAT factors, resulting in their nuclear translocation and transcriptional activity^9^.

Three members of the *Nfat* gene family, namely *Nfat1, Nfat2* and *Nfat4* are expressed in the T cell lineage and regulate many aspects of T cell functions. Although the picture is far from being complete, biochemical and gene inactivation studies have enlighted both specific, redundant or antagonistic functions of these genes in thymocyte development and mature T cell functions^11,12,13,14,15,16,17^.

In this study, we show that *Nfat1, Nfat2* and *Nfat4* are critical effectors of calcineurin in T-ALL that act mostly in a redundant fashion in regulating leukemia initiating potential and the expression of several genes/pathways that could mediate the pro-oncogenic properties of Cn/NFAT activation.

## RESULTS

### NFAT transcription factors are required for T-ALL leukemia initiating potential

We have previously shown that three NFAT factors, namely NFAT1, NFAT2 and NFAT4 are expressed and activated (dephosphorylated) in primary mouse T-ALL and diagnostic human T-ALL cases xenografted in NSG mice^7, 8^. Given the prominent role of NFAT1 in progression of solid tumors^18^, we evaluated T-ALL onset in TEL (ETV6)-JAK2 transgenic mice^19^ that are either wild-type or deficient for *Nfat1*. As reported previously, expression of the *TEL-JAK2* fusion oncogene in mouse lymphoid lineage induced T-ALL with high penetrance. We found no difference in T-ALL onset and penetrance between *Nfat1*-proficient and *Nfat1-deficient* mice (Figure 1A). Likewise, *Nfat1* inactivation failed to impact upon T-ALL onset and penetrance^20^ in the well characterized ICN1 (activated NOTCH1 allele)-induced T-ALL model (Figure 1B). These data indicate that *Nfat1* is dispensable for T-ALL development.

**Figure 1.**
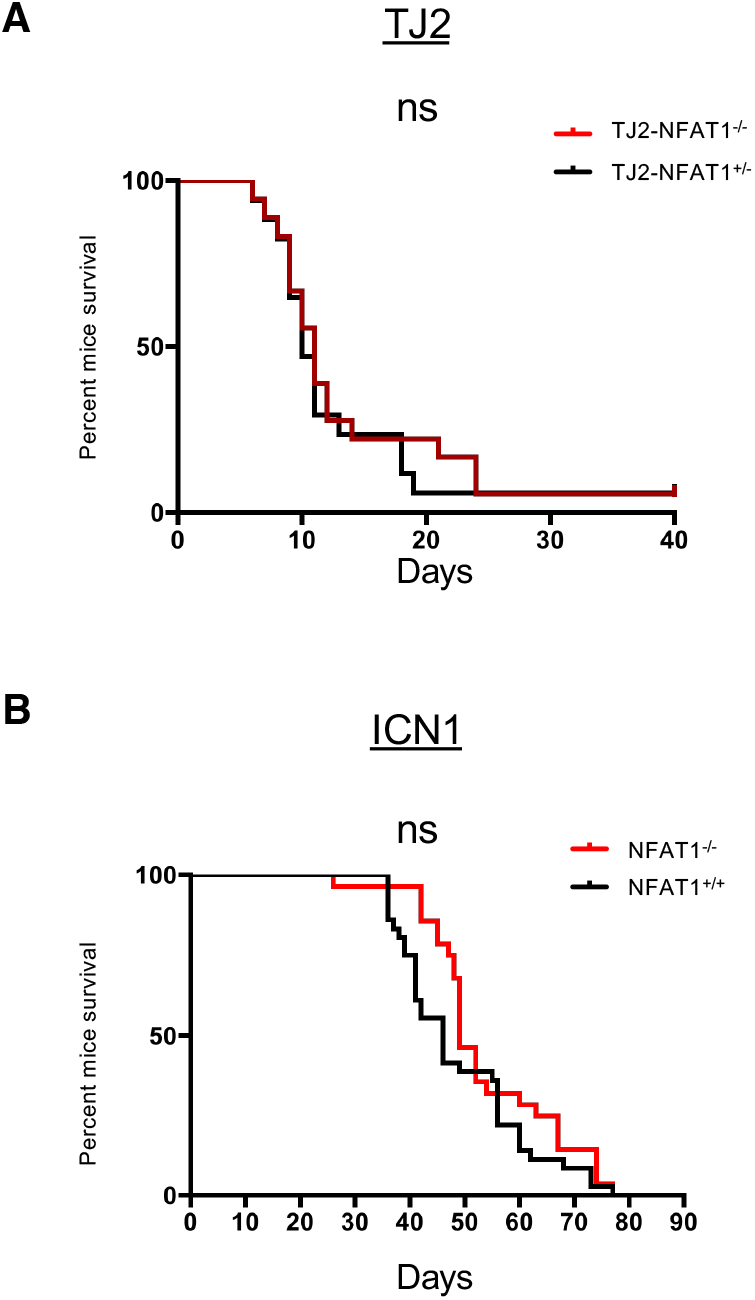
*Nfat1* gene inactivation does not affect leukemia development in TEL-JAK2 and ICN1-induced T-ALL. **(A)** TEL-JAK2 transgenic mice were crossed and backcrossed with mice inactivated for *Nfat1* to generate cohorts of TEL-JAK2^+/0^; Nfat1^+/-^ (n=17) and TEL-JAK2^+/0^; Nfat1^-/-^ (n=18) littermates. These cohorts were followed over time for T-ALL onset and mouse survival (log-rank test; ns: non significant). **(B)** Mice carrying ICN1-induced T-ALL of the indicated genotypes (*Nfat1*^+/+^, n=35; *Nfat1*^-/-^, n=28) were followed over time for T-ALL onset and recipient mice survival (log-rank test; ns: non significant).

To assess the importance of the other NFAT factors in T-ALL, we next generated *Nfat1*-deficient ICN1-driven T-ALLs carrying conditional, floxed alleles of *Nfat2* and *Nfat4* and the Rosa-Cre-ERT2 transgene (RC^T2^) (see Materiel and Methods) to ultimately compare the leukemia initiating potential (LIC activity) of NFAT-proficient and NFAT-deficient T-ALL cells. Three independent primary T-ALL (#21; #23; #24) of this genotype were thus injected into wild type secondary recipient mice. Sub-terminally leukemic mice were treated either with carrier solvent (So) or with tamoxifen (Tam) to induce Cre-mediated deletion of *Nfat2* and *Nfat4* (see Figure 2A for a scheme of the experiment). Genotyping of leukemic cells retrieved from mice 5 days later showed efficient deletion of the *Nfat4* floxed alleles (Figure 2B) associated with undetectable NFAT4 protein expression (Figure 2C) and a clear but incomplete deletion of *Nfat2* accompanied by a strong decrease in NFAT2 protein isoforms expression (Figures 2B and C). In this experimental setting in which loss of NFAT expression was experienced for about 2 days by leukemic cells (see Methods), we observed no effect on tumor burden and leukemic cell survival (Supplementary Figure 1A and B). Comparison of LIC activity of such generated *Nfat*-deficient cells with their respective *Nfat*-proficient leukemic cells was next investigated by transplantation under limit dilution conditions into new hosts and monitoring of leukemia recurrence and mouse survival (Figure 2A for a scheme). While 10^4^ (T-ALL #21) or 10^3^ (T-ALL #23; #24) *Nfat*-proficient cells were sufficient to re-initiate leukemia in all recipients, none (T-ALL #21, #24) or only 1/6 recipient mice (T-ALL #23) infused with the same number of *Nfat*-deficient T-ALL cells succumbed to leukemia (Figure 2D and Table 1). The differential phenotype between *Nfat*-proficient and *Nfat*-deficient leukemic cells did not result from a non-specific, toxic effect of tamoxifen treatment or Cre activation since the leukemia initiating potential of ICN1-driven T-ALL that are wild type for all 3 *Nfat* genes and that carry the RC^T2^ transgene is unaffected by Tam treatment (Supplementary Figure 2; Supplementary Table1). These results demonstrate that inactivation of NFAT function impairs the leukemia initiating potential of T-ALL cells.

**Figure 2.**
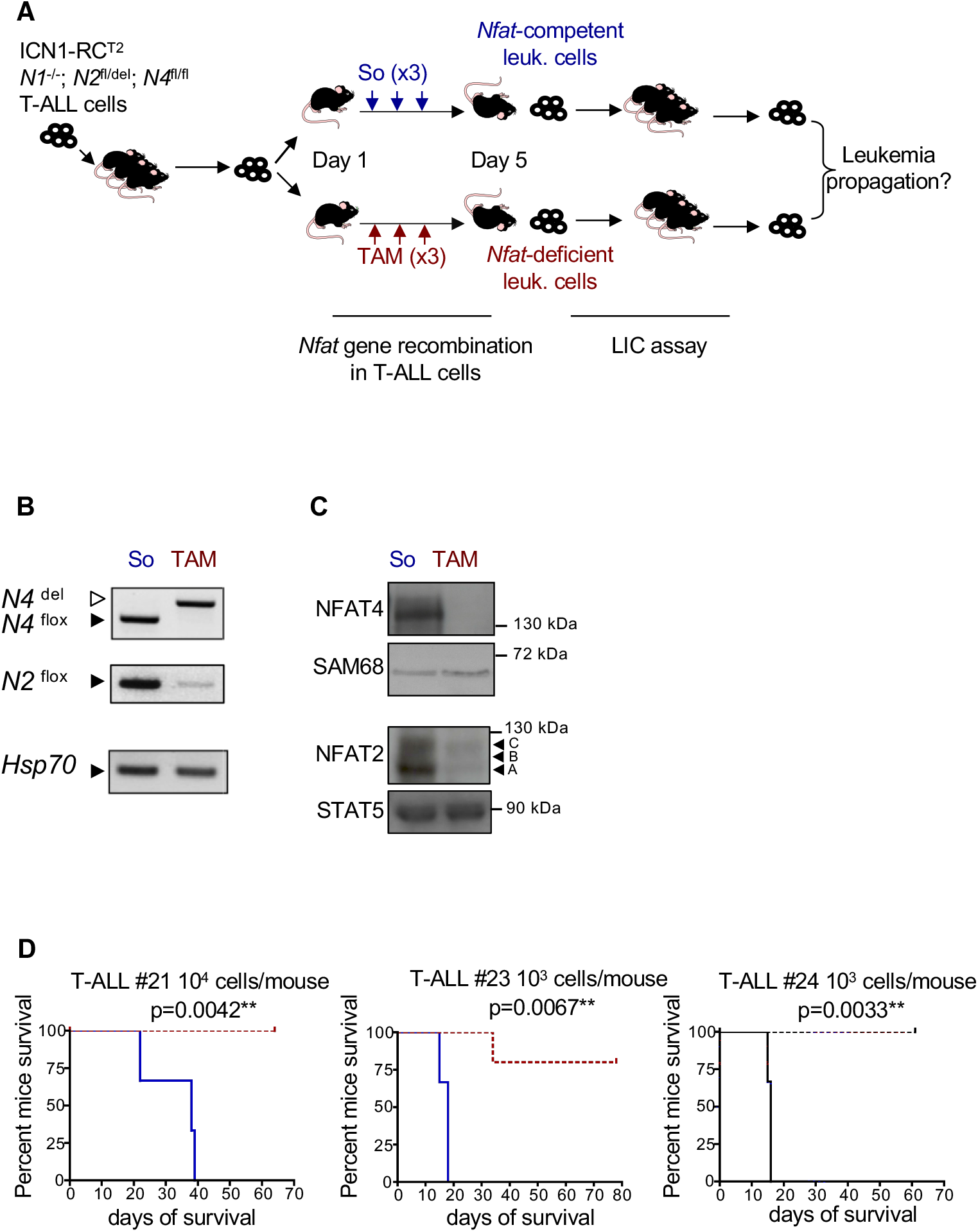
NFAT factors are required for leukemic initiating cell potential in T-ALL. **(A)** Schematic description of the experiment: Mice were i.v. injected with ICN1-induced leukemic cells obtained from 3 independent T-ALL (#21, #23, #24) of the indicated genotypes. When BM leukemia burden reached about 10-15% leukemic cells in these recipients, mice received 3 successive daily injection of either carrier solvent (So, n=3) or tamoxifen (TAM, n=6) to induce *Nfat2* and *Nfat4* floxed alleles deletion. Mice from both groups became terminally ill 2 days later and were sacrificed. *Nfat*-proficient (blue label, 1) and *Nfat*-deficient (red label, 2) leukemic cells were flow cytometry-sorted (tNGFR+ cells), genotyped for *Nfat* floxed (flox) and deleted (del) alleles and compared for their ability to re-initiate leukemia in wild-type secondary recipient mice under limit dilution conditions. *N1: Nfat1; N2: Nfat2; N4: Nfat4;* RC^T2^: Rosa-Cre-ER^T2^ transgene; ICN1: intracellular NOTCH1; LIC: leukemia initiating cell. **(B)** PCR-based genotyping for *Nfat2* and *Nfat4* floxed and deleted alleles in tNGFR+ T-ALL cells obtained from mice treated with So (1) or Tam (2), as schematized in panel A. **(C)** Western blot analysis of NFAT2 and NFAT4 expression in leukemic cells of mice treated with So (1) or Tam (2), as schematized in panel A. SAM68 and STAT5 expression are used as loading controls. Arrowheads indicate the A, B and C NFAT2 isoforms. **(D)** Kaplan-Meier survival curve of mice infused with 1×10^4^ (T-ALL #21) or 1×10^3^ (T-ALL #23; 24) *Nfat*-proficient (blue tracing) or *Nfat*-deficient (red tracing) cells as described in (A). Mice were followed overtime for T-ALL recurrence and recipient mice survival (n=3-6; log-rank test).

**Table 1.** Comparison of the leukemia initiating potential of *Nfat*-proficient and *Nfat*-deficient versions of T-ALL #21, #23, and #24, generated as schematized in Figure 1A

| Leukemia id | Treatment of donor mice | Number of cells injected in recipients | Number of injected recipients | Number of leukemic recipients (time to death, days) | Statistics | % leukemic cells in the BM ± SEM of recipients |
| --- | --- | --- | --- | --- | --- | --- |
| #21 | So | 4.10<sup>6</sup> | 3 | 3 (13 ;13 ;13) | p=0,025 | 63,9±6 |
|  | TAM | 4.10<sup>6</sup> | 3 | **3\*** (17 ;17 ;17) |  | 64,9±2,8 |
|  | So | 1.10<sup>4</sup> | 3 | 3 (22 ;39 ;38) | p=0,004 | 46,2±13,4 |
|  | TAM | 1.10<sup>4</sup> | 6 | 0 |  | NA |
|  | So | 1.10<sup>3</sup> | 3 | 1 (21) | ns | 65,9 |
|  | TAM | 1.10<sup>3</sup> | 6 | 0 |  | NA |
| #23 | So | 4.10<sup>6</sup> | 3 | 3 (13 ;14 ;13) | p=0.004 | 47,6±2,9 |
|  | TAM | 4.10<sup>6</sup> | 5 | **5\*** (20 ;20 ;25 ;22 ;29) |  | 42,9±3,9 |
|  | So | 1.10<sup>4</sup> | 3 | 3 (14 ;15 ;18) | p=0,001 | 38,1±11,1 |
|  | TAM | 1.10<sup>4</sup> | 6 | **6\*** (22 ;25 ;25 ;25 ;29 ;52) |  | 37,6±13 |
|  | So | 1.10<sup>3</sup> | 3 | 3 (18 ;15 ;18) | p=0,006 | 36,2±9,1 |
|  | TAM | 1.10<sup>3</sup> | 6 | **1\*** (34) |  | 49,3 |
| #24 | So | 4.10<sup>6</sup> | 3 | 3 (15 ;15 ;15) | p=0,004 | 58,8±5,7 |
|  | TAM | 4.10<sup>6</sup> | 6 | **6\*** (17 ;17 ;18 ;18 ;18 ;23) |  | 42,4±5,5 |
|  | So | 1.10<sup>4</sup> | 3 | 3 (15 ;15 ;15) | p=0,004 | 31,3±3 |
|  | TAM | 1.10<sup>4</sup> | 6 | **2\*** (18 ;37) |  | 39,4±8,1 |
|  | So | 1.10<sup>3</sup> | 3 | 3 (15 ;15 ;16) | p=0,003 | 9±1,6 |
|  | TAM | 1.10<sup>3</sup> | 6 | 0 |  | NA |

We next investigated whether NFAT transcription factors were involved in the control of survival, proliferation and migration of leukemic cells in these conditions, as previously observed for calcineurin. For this, ICN1; RC^T2^; *Nfat1^-/-^; Nfat2^fl/del^; Nfat4^fl/fl^* T-ALL cells were treated either with 4-hydroxytamoxifen (4OHT) to induce deletion of *Nfat2* and *Nfat4* or with the carrier solvent (Et), as control and co-cultivated with MS5 stromal cells^6,8^. *Nfat* inactivation (Figure 3A) was accompanied by impaired S phase progression as measured by BrdU pulse-labeling (Figure 3B), increased apoptosis as analyzed by caspase 3 activation (Figure 3C) and decreased cell migration as determined by time lapse videomicroscopy (Figure 3D). In contrast, ICN1; RC^T2^; *Nfat1*^+/+^; *Nfat2*^+/+^; *Nfat4*^+/+^ T-ALL cell migration, survival and proliferation were not affected by 4OHT treatment (data not shown), showing the specificity of the observed phenotypes.

**Figure 3.**
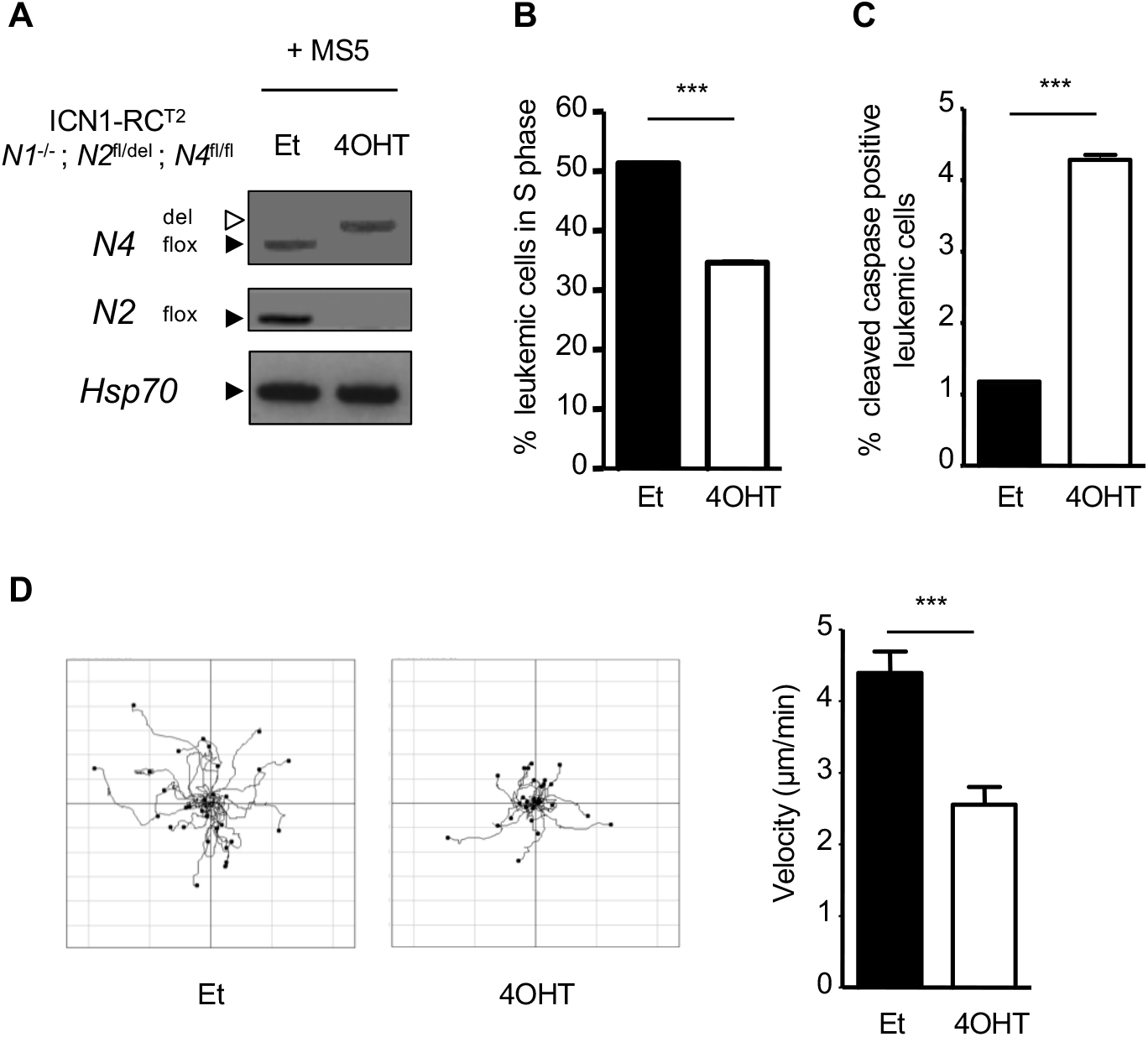
NFAT transcription factors regulate cell survival, proliferation and migration of ICN1-induced T-ALL *in vitro*. **(A)** T-ALL #21 leukemic cells were co-cultured on MS5 stromal cells and treated with solvent (ethanol, Et) or 4OHT to induce deletion of *Nfat* floxed alleles. Leukemic cells genotypes were analyzed 5 days later by PCR for the floxed (flox) and deleted (del) alleles of *Nfat4* and the floxed allele of *Nfat2.* PCR for *Hsp70* is used as control. **(B)** BrdU pulse-labeling analysis *Nfat*-proficient and *Nfat*-deficient 5 days after ethanol or 4OHT treatment, respectively. Percentage BrdU-positive leukemic cells is presented (data are represented as ± SEM; n=3; Student’s t-test; *** p<0,001). **(C)** At the same time point, *Nfat*-proficient (black) and *Nfat*-deficient (white) leukemic cells were analyzed for percentage of cells positive for cleaved caspase 3 (data are represented as mean ± SEM; n=3; Student’s t-test; *** p<0,001). **(D)** At the same time point, *Nfat*-proficient and *Nfat*-deficient leukemic cells were seeded on MS5 stromal cells and migration of individual cells (n=30) recorded for 15 minutes by time-lapse videomicroscopy. In the flower plot diagrams (left), the starting point of each track is placed at the axis origins. In the right panel, velocity (μm/min) of *Nfat*-proficient (Et) versus *Nfat*-deficient (4OHT) leukemic cell was compared (data are represented as mean ± SEM; n=3; Student’s t-test *** p<0,001).

### NFAT transcription factors have redundant functions in T-ALL

Given their redundant, agonistic or antagonistic roles during T cell development, we next investigated whether expression of individual members of the NFAT family would be sufficient to sustain leukemia initiating potential of T-ALL cells. To this end, we generated ICN1-driven tumors in which only *Nfat4* (ICN1; RC^T2^; *Nfat1*^-/-^; *Nfat2^fl/fl^*; *Nfat4*^+/+^) or only *Nfat2* (ICN1; RC^T2^; *Nfat1*^-/-^; *Nfat2*^+/+^; *Nfat4^fl/fl^*) will remain expressed following genetic inactivation of the other family members. Of note, in this experimental setting neither the loss of NFAT2 nor that of NFAT4 for about 2 days in leukemic cells (see Materiel and Methods) impacted tumor burden (Supplementary Figure 3A and B). We next analyzed the leukemia initiating potential of these cells (see Figure 4A for a scheme of the experiment). When injected under limiting dilution conditions, the LIC potential of T-ALL cells expressing only *Nfat4* was comparable to that of cells expressing both *Nfat2* and *Nfat4* (Figure 4B, left panel, compare red and blue tracings; Table 2). Of note, T-ALL cells recovered from terminally leukemic recipient mice injected with RC^T2^; *Nfat1*^-/-^; *Nfat2*^del/del^; *Nfat4*^+/+^ T-ALL cells kept their original *Nfat* genotypes (Figure 4C, left panel), indicating that the mere expression of NFAT4 is sufficient to maintain the LIC potential of T-ALL. Likewise, recipient mice injected under limit dilution conditions with leukemic cells expressing only *Nfat2* all succumbed to T-ALL, although with a slight delay as compared to mice infused with their respective control (Figure 4B, right panel compare red and blue tracings; Table 2). Leukemic cells recovered from terminally-leukemic recipients injected with RC^T2^; *Nfat1*^-/-^; *Nfat2*^+/+^; *Nfat4*^del/del^ T-ALL kept their original *Nfat* genotypes (Figure 4C; right panels), showing that LIC activity was effectively driven by NFAT2. Taken together, these experiments show that NFAT factors play an essential and redundant role in the leukemia initiating potential of T-ALL cells.

**Figure 4.**
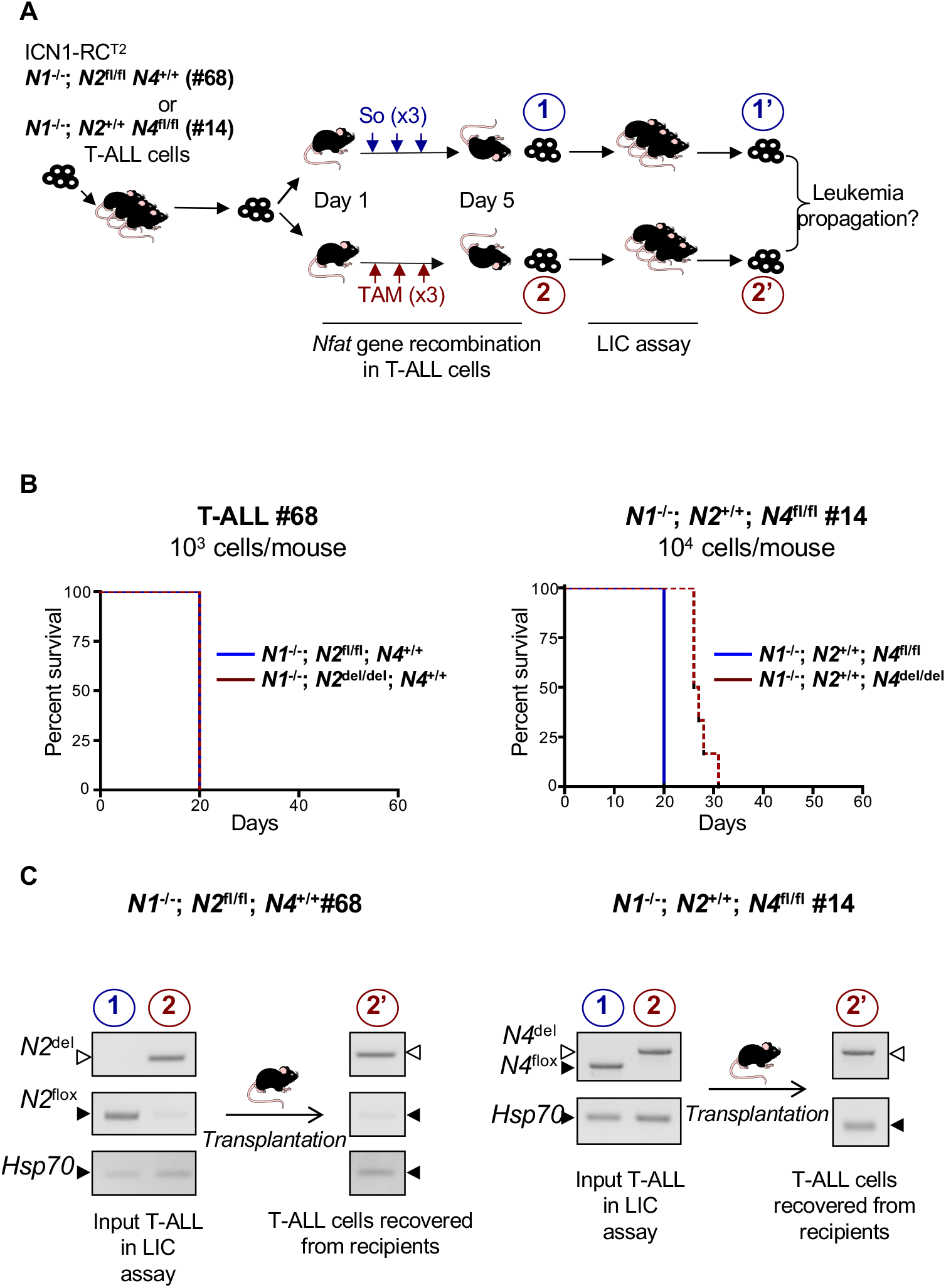
Functional redundancy of NFAT factors in T-ALL. **(A)** Schematic description of the experiment: mice were injected with leukemic cells obtained from either primary T-ALL #68 (ICN1; RCT2; *Nfat1*^-/-^; *Nfat2*^fl/fl^; *Nfat4*^+/+^) or #14 (ICN1; RCT2; *Nfat1*^-/-^; *Nfat2*^+/+^; *Nfat4*^fl/fl^), that carry wild type alleles of *Nfat4* or *Nfat2*, respectively. When BM leukemia burden reached about 10-15% T-ALL cells in these recipients, mice received 3 successive daily injection of either carrier solvent (So, n=3) or Tamoxifen (TAM, n=6) to induce *Nfat2* (T-ALL #68) or *Nfat4* (T-ALL #14) floxed alleles deletion thus resulting in T-ALLs relying upon only *Nfat4 (T-ALL #68) or only Nfat 2 (T-ALL #14)*. Mice from all groups became terminally leukemic 2 days later and *Nfat*-proficient (blue label, 1) or *Nfat*-defloxed (red label, 2) cells were flow cytometry-sorted and compared for their ability to re-initiate leukemia in secondary recipient mice under limit dilution conditions. Flow cytometry-sorted leukemic cells obtained from donor mice (1, 2) and retrieved from terminally ill recipients (1’, 2’) were genotyped for *Nfat* floxed and deleted alleles. **(B)** Left panel: Kaplan-Meier survival curves of mice transplanted with T-ALL cells #68 expressing only *Nfat4* (red tracing) or co-expressing *Nfat2* and *Nfat4* (blue tracing). Right panel: Kaplan-Meier survival curves of mice transplanted with T-ALL cells #14 expressing only *Nfat2* (red tracing) or co-expressing *Nfat2* and *Nfat4* (blue tracing). **(C)** Left panels: PCR genotyping of *Nfat2* floxed and deleted alleles in input leukemic cells from T-ALL #68 (1 and 2; see schematic in A) and in leukemic cells recovered from one representative secondary recipient injected T-ALL cells (T-ALL #68, 2’). Right panels: PCR genotyping of Nfat4 floxed and deleted alleles in input leukemic cells from T-ALL #14 (1 and 2; see schematic in A) and in leukemic cells recovered from one representative secondary recipient injected with T-ALL #14 (2’). PCR for *Hsp70* is used as control.

**Table 2.** Comparison of the leukemia initiating potential of Nfat-floxed and Nfat-defloxed versions of T-ALL #68 and T-ALL #14, generated as described in Figure 4A.

| Leukemia id | Treatment of donor mice | Number of cells injected in recipients | Number of injected recipients | Number of engrafted recipients (time to death, days) | Statistics | % leukemic cells in the BM ± SEM of recipients |
| --- | --- | --- | --- | --- | --- | --- |
| N1<sup>-/-</sup>;N2<sup>fl/fl</sup>;N4<sup>+/+</sup><br>#68 | So | 1.10<sup>6</sup> | 3 | 3 (10 ;10 ;10) | ns | 32,6±2,2 |
| | TAM | 1.10<sup>6</sup> | 6 | 6 (10 ;10 ;10 ;10 ;10 ;10) | | 26,6±1 |
| | So | 1.10<sup>4</sup> | 3 | 3 (16 ;16 ;16) | ns | 35,1±1,4 |
| | TAM | 1.10<sup>4</sup> | 6 | 6 (16 ;16 ;16 ;17 ;17 ;17) | | 30,9±3,2 |
| | So | 1.10<sup>3</sup> | 3 | 3 (20 ;20 ;20) | ns | 34,2±3,4 |
| | TAM | 1.10<sup>3</sup> | 5 | 5 (20 ;20 ;20 ;20 ;20) | | 30,7±1,5 |
| | So | 1.10<sup>2</sup> | 3 | 2 (20 ;31) | ns | 38,8±5,1 |
| | TAM | 1.10<sup>2</sup> | 6 | 4 (20 ;20 ;29 ;46) | | 28,3±2,1 |
| N1<sup>-/-</sup>;N2<sup>+/+</sup>;N4<sup>fl/fl</sup><br>#14 | So | 1.10<sup>6</sup> | 3 | 3 (17 ;17 ;17) | p=0,005 | 72,7±3,6 |
| | TAM | 1.10<sup>6</sup> | 6 | 6 (18 ;18 ;20 ;20 ;21 ;21) | | 62,2±12,0 |
| | So | 1.10<sup>4</sup> | 3 | 3 (20 ;20 ;20) | p=0,005 | 61,2±6,4 |
| | TAM | 1.10<sup>4</sup> | 6 | 6 (26 ;26 ;26 ;27 ;28 ;31) | | 61,4±10,4 |
| | So | 1.10<sup>3</sup> | 3 | 3 (21 ;21 ;21) | p=0,05 | 66,2±3,4 |
| | TAM | 1.10<sup>3</sup> | 6 | 5 (27 ;27 ;35 ;38 ;38) | | 51,8±0,8 |
|  | So | 1.10<sup>2</sup> | 3 | 1 (28) | ns | 70,7 |
|  | TAM | 1.10<sup>2</sup> | 6 | 1 (60) |  | 8,2 |

### NFAT factors act downstream of Cn

Our results show that NFAT deficiency essentially phenocopies Cn deficiency in T-ALL^8^, suggesting NFAT to be central effectors of Cn in T cell leukemogenesis. To investigate this hypothesis, we analyzed whether expression of constitutively active NFAT mutants could compensate the cellular phenotypes linked to Cn inactivation. We used the constitutive mutant NFAT1[2+5+8]^21^ (named NFAT1* thereafter) in which most serine residues targeted by NFAT kinases in the SRR1, SP2 and SP3 motifs were mutated into alanine, thus mimicking calcineurin-induced dephosphorylation and the same mutant carrying in addition a SV40 nuclear localization signal (NLS) motif at its C-terminus (NFAT1*-NLS), further enhancing its nuclear accumulation and transcriptional activity^21^. ICN1; RC^T2^; CnB1^fl/fl^ T-ALL cells^8^, were retrovirally transduced at the same m.o.i. either with the GFP control vector (MIG), or MIG vectors encoding HA-tagged version of either CnB1, or with the constitutive NFAT1 mutants and concomitantly treated with 4OHT to induce *CnB1 (PPP3R1)* gene deletion and calcineurin inactivation^8^ (Figure 5A for a schematic representation of the experiment). Leukemic cells were then co-cultured with MS5 stromal cells and followed over time. As shown in Figure 5B and as previously reported, 4-OHT treatment induced efficient *CnB1* gene deletion (Figure 5B) and resulted in a strong arrest in T-ALL expansion (Figure 5C), with the few cells found under these co-culture conditions resulting from the survival of a minor population of cells that escaped full *CnB1* gene deletion, detectable by PCR (Figure 5D, MIG lanes). As expected, expression of exogenous CnB1 restored the ability of T-ALL cells to survive and proliferate (Figure 5C, left panel, blue tracing) concomitant with the rapid amplification of GFP+ cells (Figure 5C, right panel, blue tracing), that retain the parental CnB^del/del^ genotype (Figure 5D).

**Figure 5.**
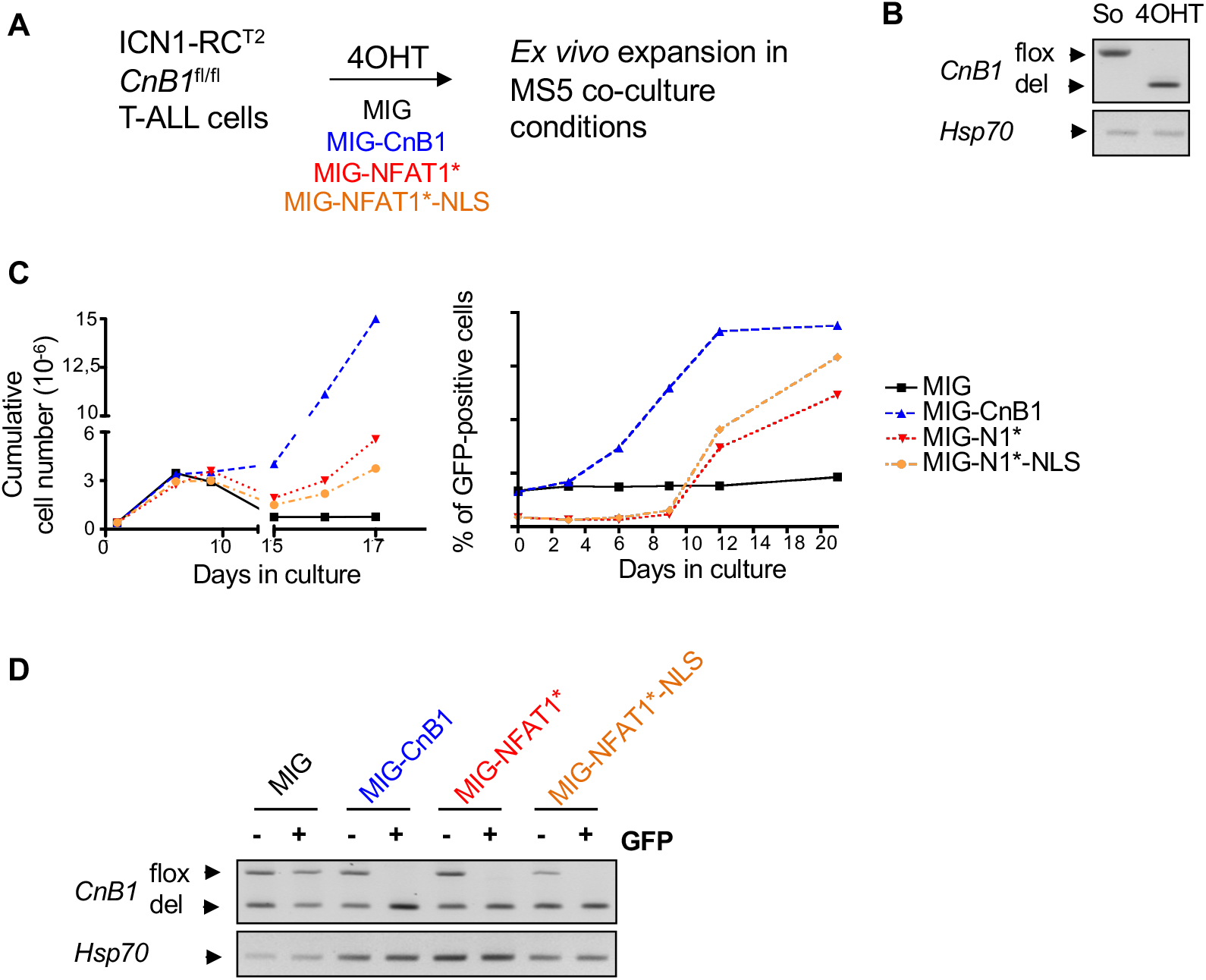
NFAT factors are major downstream effectors of calcineurin in T-ALL. **(A)** Schematic of the experiment: leukemic cells obtained from an ICN1-induced T-ALL carrying 2 floxed alleles of *CnB1* (T-ALL #3) were retrovirally transduced with MIG vectors encoding either HA-tagged CnB1, or HA-tagged NFAT1*, or HA-tagged NFAT1*-NLS, or the control vector without insert (MIG), co-cultured on MS5 stromal cells and immediately treated with 4OHT to delete *CnB1* floxed alleles. **(B)** PCR genotyping analysis for the floxed and deleted alleles of *CnB1* in cultured cells 2 days after 4OHT treatment. PCR for *Hsp70* is used as control. **(C)** Left panel: Expansion of CnB1-deleted leukemic cells transduced with MIG (black squares), MIG-CnB1 (blue triangle), MIG-NFAT1* (red triangles), MIG-NFAT1*-NLS (orange); expansion over time is reported as the cumulative cell numbers over 17 days in MS5 co-cultures. Right panel: in the same co-cultures, enrichment in GFP+, transduced cells, was followed over time. This experiment is representative of three independent experiments. **(D)** PCR genotyping analysis for the floxed and deleted alleles of *CnB1* in the GFP^+^ (transduced) and GFP^-^ (non-transduced) fractions of flow-cytometry sorted leukemic cells from the respective co-cultures at day 17. PCR for *Hsp70* is used as control.

Interestingly, both NFAT1* and NFAT1*-NLS rescued the survival/proliferation defect of CnB1-deficient T-ALL cells (Figure 5C red and orange tracing, respectively) albeit less efficiently as compared to exogenous CnB1. Cn-deficient T-ALL cells transduced with either of the NFAT1* vectors (GFP+) maintained the original CnB1-deleted genotype unlike the remaining fraction of non-transduced cells (GFP-) found in these cultures (Figure 5D). We conclude from these experiments that constitutive NFAT activity can restore the survival/proliferative properties of Cn-deficient T-ALL cells, indicating that NFAT are major downstream effectors of Cn in T-ALL.

### NFAT-dependent transcriptome in T-ALL

To gain insight into the molecular basis of NFAT oncogenic properties, we compared the transcriptome of *Nfat*-proficient and *Nfat*-deficient leukemic cells, obtained as described in Figure 1A, using 3 independent ICN; RC^T2^; *Nfat1^-/-^; Nfat2^fl/fl^; Nfat4^fl/fl^* T-ALL. Hierarchical clustering analysis clearly distinguished NFAT-proficient from NFAT-deficient cells (Figure 6A) with 343 probe sets being significantly deregulated (FC>1,5 and p<0,05, Supplementary Table 2). IPA analysis of this signature did not highlight deregulation of a specific pathway (data not shown). However, we noticed the up-regulation of a number of genes encoding proteins that are either physiological regulators in T cells, e.g. B- and T-cell attenuator (*Btla*), an inhibitory receptor of T cell signaling belonging to the CD28 superfamily; *Gimap7*, a member of the immunity associated GTPases, regulators of lymphocyte survival and homeostasis; *Nr4a1*, a nuclear protein involved in thymocyte clonal deletion during negative selection; *Irf4*, a transcriptional regulator involved in multiple aspects of T cell development. *Nfat* inactivation also deregulated genes implicated in cell cycle regulation (e.g. up-regulation of *Cdkn1A, Gadd45b;* down-regulation of *Evi5*) (Supplementary Table 2). Of note, comparison of the NFAT-regulated transcriptome with that regulated by calcineurin in ICN1 T-ALL^8^ identified 96 commonly deregulated probe sets (**Supplementary Table 3**). Gene enrichment analysis revealed a robust overlap between the Calcineurin and Nfat-dependent transcriptome, particularly amongst the upregulated processes (Figure 6B; Supplementary Table 3). We identified a node of particular interest, accounting for proteins implicated in the regulation of cell proliferation and survival (Figure 6C). RT-PCR experiments performed in independent T-ALL confirmed that both *Cdkn1a* and *Nr4a1* were up-regulated in response to either NFAT or CnB-1 inactivation (Figure 6D), validating at the molecular level the resemblance of CnB1 and NFAT loss of function phenotypes.

**Figure 6.**
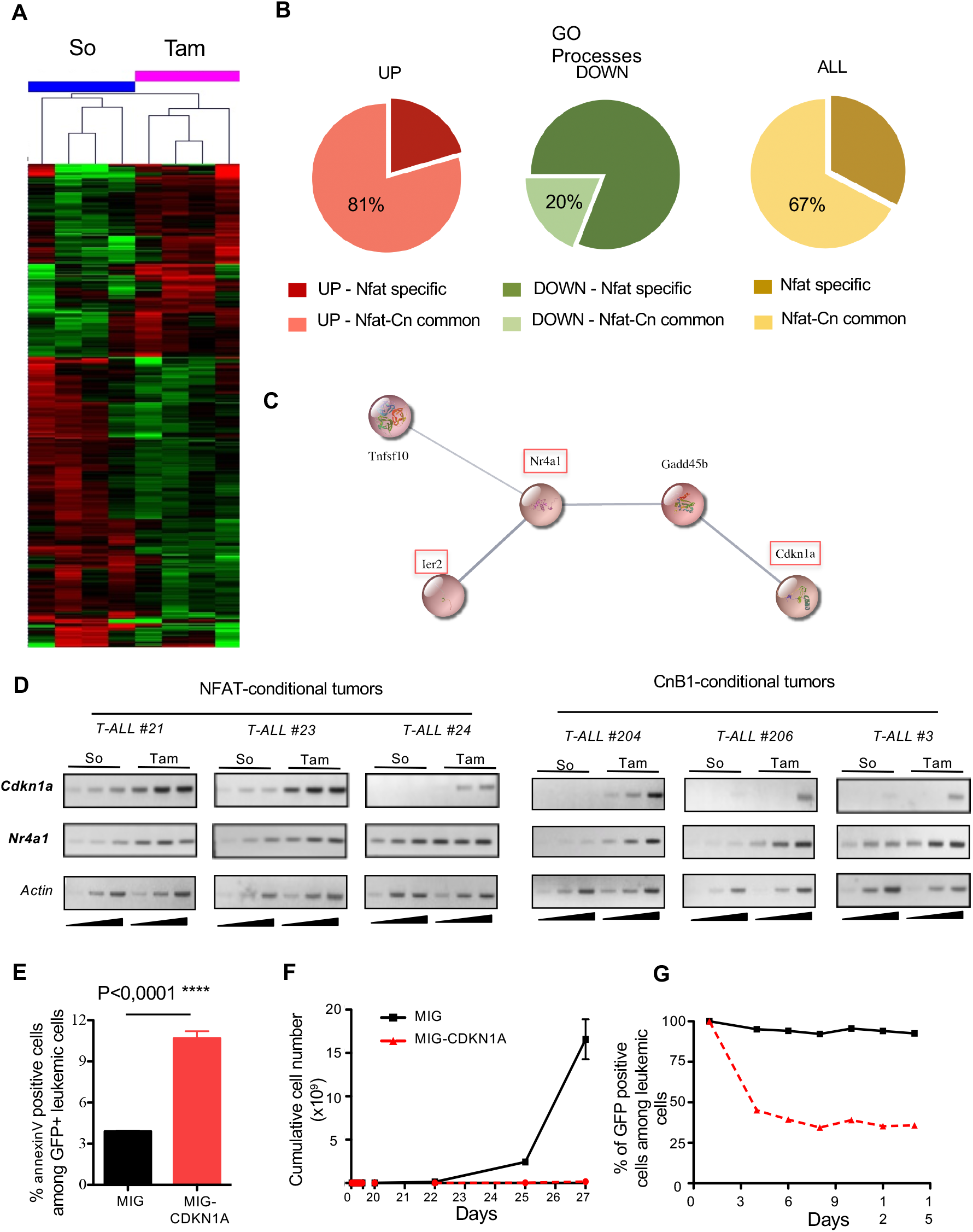
NFAT-dependent transcriptome analysis in ICN1-induced T-ALL. **(A)** Global gene expression analyses of *Nfat*-proficient and *Nfat*-deficient leukemic cells obtained from T-ALL #21, #24, #23 (2 independent experiments) and sorted by flow cytometry from solvent (So)- or Tam-treated mice generated as described in Fig2A. Hierarchical clustering of the different leukemias (top legends) in their *Nfat*-competent (So) and *Nfat-deficient* (Tam) versions was performed using a fold change ≥0,9 with a p value <0,05. The heatmap representation highlights up-regulated genes in red and down-regulated genes in green. **(B)** Pie charts showing the percentages of commonly deregulated GO processes in the Cn-dependent and *Nfat*-dependent transcriptome. **(C)** Predicted protein interaction map retrieved from the analysis of significantly upregulated genes in the T-ALL *Nfat*-dependent transcriptome. Orange boxes point to genes commonly deregulated in both NFAT- and Cn-dependent transcriptomes **(D)** *Nfat*-proficient, *Nfat*-deficient, *CnB1*-proficient and *CnB1*-deficient versions of the indicated T-ALL were sorted by flow cytometry and analyzed by semi-quantitative RT-PCR for the expression of *Cdkn1a* and *Nr4a1.* RT-PCR for expression of *β-actin* is used as control **(E)** Leukemic cells from T-ALL #3 were transduced with MIG vectors encoding CDKN1A or the control MIG vector without insert. Leukemic cells survival was analyzed by Annexin V staining at day 4 in co-cultures of the indicated leukemic cells with MS5 stromal cell (data are represented as ± SEM; n=3; Student’s t-test). **(F)** Expansion over time in MS5 co-cultures of leukemic cells transduced with the MIG and MIG-CDKN1A vectors described in E. **(G)** Percentage of transduced (GFP+) leukemic cells in co-cultures described in (F) was followed by flow cytometry.

To gain insight into the functional relevance of this deregulated node, we next investigated the consequences of CDKN1a overexpression on T-ALL expansion. For this, leukemic cells were retrovirally-transduced with the MIG control vector or MIG vector encoding p21^CDKN1a^. Western blot analysis confirmed p21^CDKN1a^ overexpression as compared to MIG-transduced cells (Supplementary Figure 4). Enforced expression of p21^CDKN1a^ resulted in induction of apoptosis and impaired leukemic cell expansion in MS5 co-cultures (Figure 6E and F), with transduced cells (GFP+) rapidly being counterselected (Figure 6G). Similar results were obtained upon enforced expression of Nr4a1 while enforced expression of Gadd45b was without effect (data not shown). Taken together, these results indicate an essential role of NFAT-regulated genes in orchestrating important leukemic phenotypes in T-ALL.

## DISCUSSION

We previously demonstrated through pharmacological and genetic approaches that calcineurin is important to T-ALL maintenance *in vivo* and *ex vivo* and critical to their leukemia propagating potential^7, 8^. We now demonstrate that the three NFAT factors activated in T-ALL, namely NFAT1, 2 and 4 are also essential to the survival/proliferation/migration properties and leukemia propagating potential of T-ALL. We found that only the concomitant inactivation of all three NFAT factors results in impaired LIC activity in T-ALL, as evidenced by the fact that expression of either NFAT2 alone or NFAT4 alone was sufficient to maintain this activity. This demonstrates clear functional redundancy for these factors in T-ALL. Since expression of a constitutive mutant of NFAT1 that mimics NFAT1 dephosphorylated, active state restores the survival and proliferation properties of calcineurin-deficient leukemic cells and LIC potential (data not shown), our results highlight the central function of NFAT downstream of calcineurin activation in T-ALL.

The redundant function of NFAT factors in T-ALL contrasts with previous studies interrogating of NFAT involvement in other oncogenic settings. For example, while several reports have shown that constitutively activated mutant of the a isoform of NFAT2 (the so-called short NFAT2 isoform) transforms mouse 3T3L1 adipocytes and NIH3T3 fibroblasts *in vitro*^18, 22, 23^, a constitutively activated phosphorylation mutant of NFAT1 (named NFAT1* NLS in the present study) was unable to recapitulate those effects, and even inhibited cell transformation by constitutively active NFAT2 or the Ha-RAS oncoprotein^23^. Likewise, NFAT1 *ex vivo* studies have suggested that NFAT1 has a non-redundant function in the invasive properties of breast carcinoma cell lines^24^ while, in mice, the tumorigenic/metastatic potential of mammary tumor cells rather depends upon non redundant functions of NFAT1 and NFAT2^25^. At the molecular level, NFAT2, but not NFAT1, is recruited to the *c-MYC* gene promoter leading to c-MYC over-expression, which is required for tumor maintenance together with the deregulation of other survival genes in aggressive B cell lymphomas^26^.

The *Nfat1* gene has been shown to restrain the proliferation of naive T cells *in vivo*^27, 28^ and to exert tumor suppressive functions in the mouse B cell lineage^29^. Our data show that loss of NFAT1 function does not affect leukemia onset and outcome in T-ALL induced in mice by activated NOTCH1 (ICN1) or activated JAK2 (TEL/ETV6-JAK2), indicating lack of tumor suppressive function of *Nfat1* in early T cell progenitors. We found that although expression of the constitutive mutant NFAT1*-NLS rescues survival/proliferation of CnB1-deficient T-ALL cells, it is deleterious to the survival/proliferation of CnB1-proficient ICN1 T-ALL cells *ex vivo* and *in vivo* (data not shown). This indicates that NFAT activity must be finely regulated to sustain its pro-leukemic activity in T-ALL. In line with this, fine-tuned regulation of NFAT activity is also recognized as being essential under physiological conditions for instance during thymocytes development^30^.

Comparison of the NFAT-dependent transcriptome with that associated with calcineurin inactivation in ICN1-induced T-ALL^8^ identifies a common signature, with 67% of NFAT regulated genes being also Cn-regulated. This Cn/NFAT signature in T-ALL differs from the NFAT-dependent signature characteristic of the TCR-dependent activation of peripheral T cells as no difference in expression in e.g. the genes encoding IL2, IL3, IL4, IL5, IL13, IFNg, GM-CSF was found upon *CnB1* or *Nfat* deletion in T-ALL cells. Besides T cell differentiation and activation, NFAT is also involved in exhaustion of activated T cells to limit or constrain the immune response^16, 31^. Although a trend was observed for inhibition of the expression of *Pdcd1, Lag-3 and Ctla4* (genes encoding receptors involved in NFAT-mediated exhaustion) in *Nfat*-deficient T-ALL, their differential expression in our global transcriptomic analyses did not reach statistical significance.

Instead we found NFAT to impact T-ALL maintenance through deregulation of genes with demonstrated inhibitory properties on survival, cell cycle progression of normal T cell progenitors. *Cdkn1a*, a gene recurrently altered in T-ALL diagnostic samples through promoter methylation^32^ and *Nr4a1*, a gene involved in clonal deletion of self-reactive T cells during thymic negative selection and in activated T cell exhaustion^33, 34^ are up-regulated upon either *CnB1* or *Nfat* inactivation and contribute to ICN1 T-ALL maintenance. Available evidence indicates that these genes can be regulated by NFAT factors independently of their binding to specific DNA sequences but rather through protein-protein interactions with other transcription factors^35, 36^. We also found Cn/NFAT-dependent expression of *Tox* in ICN1-induced T-ALL*. Tox* encodes a protein that facilitates genomic instability in T-ALL and essential for T-ALL cell lines survival/ proliferation and for *in vivo* maintenance^37^. Because constitutive expression of CDKN1a (this study), NR4A1 (data not shown) or TOX knockdown^37^ are sufficient to partially mimic the deleterious phenotypes induced upon NFAT or calcineurin deletion, this suggests that the Cn and NFAT commonly deregulated genes are central mediators of the pro-oncogenic properties of Cn/NFAT pathway.

Besides these transcriptionally regulated candidates, we also identified CXCR4 cell surface expression being commonly modulated by Cn and NFAT. Since NFAT-deficient cells also present reduced CXCR4 cell surface expression level (data not shown), this defect could explain the migration default and impaired leukemia inducing potential as demonstrated for Cn-deficient leukemic cells^6^, reinforcing a major role for Cn/NFAT axis in T-ALL.

Not surprisingly, we found the number of genes regulated by calcineurin^8^ to be broader than that dependent upon NFAT (this study), indicating that in addition to NFAT, calcineurin likely acts through other effectors to enforce leukemia inducing potential in T-ALL. In line with this hypothesis, several new Cn interacting proteins were recently identified, the inactivation of which could synergize with Cn inhibition to impair T-ALL expansion^38, 39^. We also noticed a number of genes specifically deregulated upon *Nfat* deletion, but not upon *CnB1* deletion, indicating that NFAT activity can be critically regulated by upstream signaling pathways in addition to their Cn-mediated nuclear translocation. In normal early T cell progenitors, the IL7/IL7R/JAK3 signaling pathway directly regulates NFAT2 through phosphorylation on a tyrosine residue in its regulatory domain^11^. Moreover, NFAT2 activity was also identified as a target of PIM kinases independent of calcineurin activation^40^. High PIM1 expression is a biomarker in T-ALL cases with JAK/STAT activation and response of leukemic cells to endogenous IL7^41, 42^, with PIM targeting cooperating with chemotherapy to promote leukemic mice survival in T-ALL PDXs^42^. Nevertheless, we did not find NFAT activity to be modulated by IL7 in T-ALL cells (data not shown), leaving space to speculate on new NFAT regulators in this context.

While T-ALL is a highly heterogeneous disease, patient treatment relies mostly on general chemotherapeutic regimens with 20% and 50% of pediatric and adult cases relapsing respectively. Targeted therapies are thus awaited for this pathology. Targeting calcineurin using CsA or FK506 could be a therapeutical option in T-ALL, yet these drugs are associated with induction of secondary cancer^43^ and ill-characterized off-target effects that limit their usefulness. The identification of downstream effectors of calcineurin as reported here may thus open novel pre-clinical investigations paths. Inhibitors of NFAT have been developed (INCA1, 2 and 6, JapA, MA242, compound 10), that act by preventing Cn/NFAT interaction^44^, promoting NFAT degradation^45^ or interfering with NFAT binding partners^46^. Given the essential role of NFAT in T-ALL, it would be interesting to analyze in the future whether these compounds, represent valid therapeutic alternatives to overcome treatment limitations in T-ALL.

## MATERIELS AND METHODS

### Analysis of T-all mouse models

Mice carrying null (-) alleles of *Nfat1*^27^, the floxed (flox) and deleted (del) alleles of *Nfat2* (a generous gift of Dr. A. Rao) and *Nfat4*^12^ (a generous gift of Dr. GR Crabtree) and the Rosa-Cre-ER^T2^ (RC^T2^) transgene^47^ were maintained on a C57BL/6 genetic background. Mice were crossed to generate the following compound mice: RC^T2^; *Nfat1*^-/-^; *Nfat2*^flox/del^; *Nfat4*^flox/flox^, or RC^T2^; *Nfat1*^-/-^; *Nfat2*^flox/flox^; Nfat4^+/+^ or RC^T2^-*Nfat1*^-/-^;*Nfat2*^+/+^; *Nfat4*^flox/flox^. Primary ICN1 (intracellular NOTCH1 domain)-induced T-ALL were obtained following retroviral-mediated gene transfer of bone marrow (BM) cells from the respective mice using either MIN-ICN1 or MIG-ICN1, which also encode either a truncated human NGFR from an IRES-tNGFR cassette or eGFP, respectively, as described^8^. T-ALL of the different genotypes emerged within 4-6 weeks and leukemic cells (tNGFR^+^ or GFP^+^) were collected from BM for further studies. To study *Nfat* function in T-ALL, leukemic cells of the indicated genotypes were intravenously (i.v) infused into secondary wild-type mice, conditions that result in synchronous leukemia engraftment in recipients. Engraftment was followed by sacrificing mice at regular time intervals and measuring % of tNGFR^+^ (GFP^+^) leukemic cells in BM. Cre activation was induced in leukemic cells by Tamoxifen (Tam) administration (Sigma-Aldrich; 1mg/mouse, three times at 24 hours intervals). Carrier solvent (corn oil, So) was used as control. Unless otherwise stated, Tam administration was started when leukemic burden reached 10-15% T-ALL cells in BM (usually 10-14 days after leukemic cells infusion). In these conditions, Cre-mediated loss of NFATs activity is experienced by leukemic cells 2 days later^8^. Mice were sacrificed when becoming moribund (5 days after the start of treatment). To study NFAT factors function in T-ALL leukemia initiating potential, NFAT-proficient and NFAT-deficient leukemic cells obtained from carrier solvent or Tam-treated mice were injected i.v at different doses (from 10^6^ to 10^3^ leukemic cells/mouse) into wild-type, syngeneic recipient mice. Time to death and leukemia burden, as analyzed by flow cytometry as % FSC large/tNGFR^+^ (or GFP^+^) leukemic blasts in BM, spleen and liver, were recorded. Mice were maintained under specific pathogen-free conditions in the animal facility of the Institut Curie. Experiments were carried out in accordance with the European Union and French National Committee recommendations, under agreement APAFIS #7393-2016102810475144-v1

### Microarray analysis

Total RNA was isolated from flow-cytometry-sorted leukemic cells (tNGFR+) 5 days following treatment with either tamoxifen or carrier solvent, using the Rneasy kit (QIAGEN). cRNA synthesis and hybridization of Mouse GeneChip® 430 2.0 arrays (Affymetrix) were according to the manufacturer’s instructions, as described (http://www.microarrays.u-strasbg.fr). A paired Student’s t-test was performed to compare gene intensities in the different biological replicates. Genes were considered significantly regulated when fold-change was ≥ 1.9 and *p* value ≤ 0.05. Significantly deregulated genes from our dataset (NFAT) and from (Cn)^8^ were used to perform pathways and processes enrichment analysis via the online STRING platform. The percentage of shared significantly upregulated, down regulated and overall deregulated processes are shown. Using a spring model via the STRING platform, we generated a network of predicted associated upregulated proteins.

### Flow cytometry, apoptosis, proliferation and migration analyses

Surface staining of tNGFR leukemic cells was performed with PE-conjugated anti-human CD271 antibody (BD Biosciences). Apoptosis assays were carried out as described^8^, using PE-conjugated-anti-active Caspase-3 antibody (BD Biosciences). BrdU incorporation assays were performed as described^6^, using APC-conjugated anti-BrdU antibody (BD Biosciences). Flow cytometry acquisitions were carried out on a FACSCalibur™ analyzer (BD Biosciences) equipped to detect 4 fluorescent parameters with the assistance of BD CellQuest Software (BD Biosciences) and data were analyzed with FlowJo Software (Tree Star). ICN1 leukemic cells were sorted on a FACSAria^™^III (BD Biosciences) cell sorter on the basis of tNGFR and/or GFP expression with the assistance of BD FACSDiva Software (BD Biosciences). Migration analyses were performed by videomicroscopy, as described^6^.

### Statistics

Statistical analyses were performed with GraphPad Prism (version 6.0; GraphPad Software, Inc.). The data are expressed as mean ± standard deviation (s.d) of n = 3 or more determinations. Unpaired two-tailed Student’s *t* tests were used to analyze experimental data between two groups. For three or more groups, a one-way ANOVA was performed using Tukey’s test. Overall survival of mice infused with ICN1-induced T-ALL was calculated according to the Kaplan-Meier method. Log-rank test was used to analyze survival curves comparisons. Differences were considered statistically significant at *p* < .05 (*),*p* < .01 (**) or *p* < .001 (***).

## AKNOWLEGMENTS

This work was supported by funds from Institut Curie, Centre National de la Recherche Scientifique (CNRS), Institut National de la Santé et de la Recherche Médicale (INSERM), Ligue contre le Cancer and Fondation ARC pour la recherche contre le cancer. CC, DP and SG were supported by pre-doctoral fellowships from the University Paris-Diderot and Ligue Nationale Contre le Cancer. The authors thank E Belloir, C Alberti and C Roulle (Institut Curie animal facility) for expert technical assistance.

## COMPETING INTERESTS

CTQ and JG have research agreements with Servier and Autolus Ltd. The remaining authors declare no competing financial interests.

## SUPPLEMENTAL FIGURES AND TABLES

**Supplementary Figure 1.**
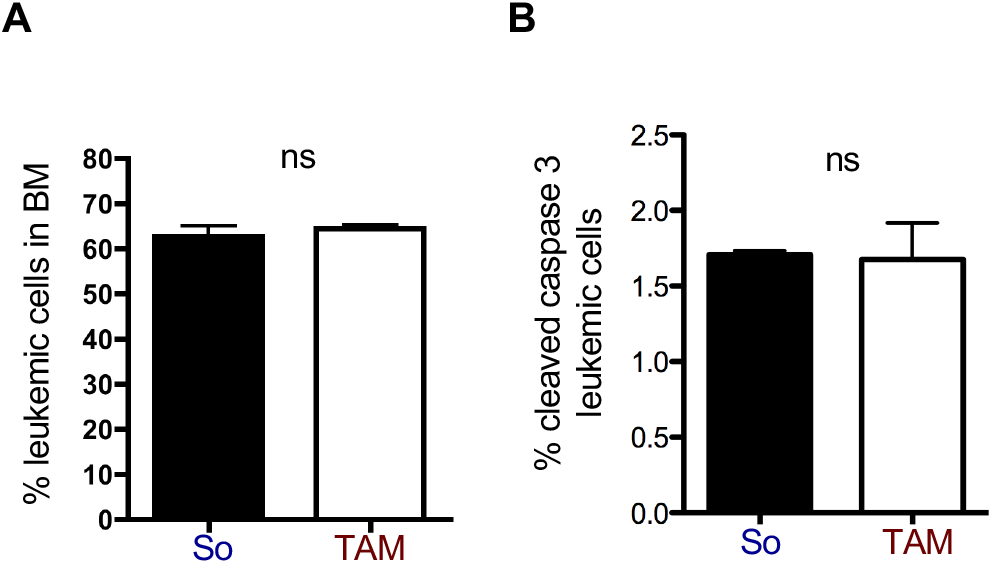
Analysis of the consequences of short-term *Nfat* inactivation in ICN1-induced T-ALL. The *Nfat*-proficient and *Nfat*-deficient versions of T-ALL #21 were generated as described in Fig1A. Mice were sacrificed when terminally ill, 2 days after the end of So and Tam treatments (1 and 2 in Fig1A). **(A)** Leukemic burden (% tNGFR+ cells) in BM was analyzed by flow cytometry at the time mice were killed (data are represented as ± SEM; n=3; Student’s t-test; ns: non significant). **(B)** Apoptosis in leukemic cells described in (A) was analyzed by measuring caspase 3 activation by flow cytometry (data are represented as ± SEM; n=3; Student’s t-test; ns: non significant).

**Supplementary Figure 2.**
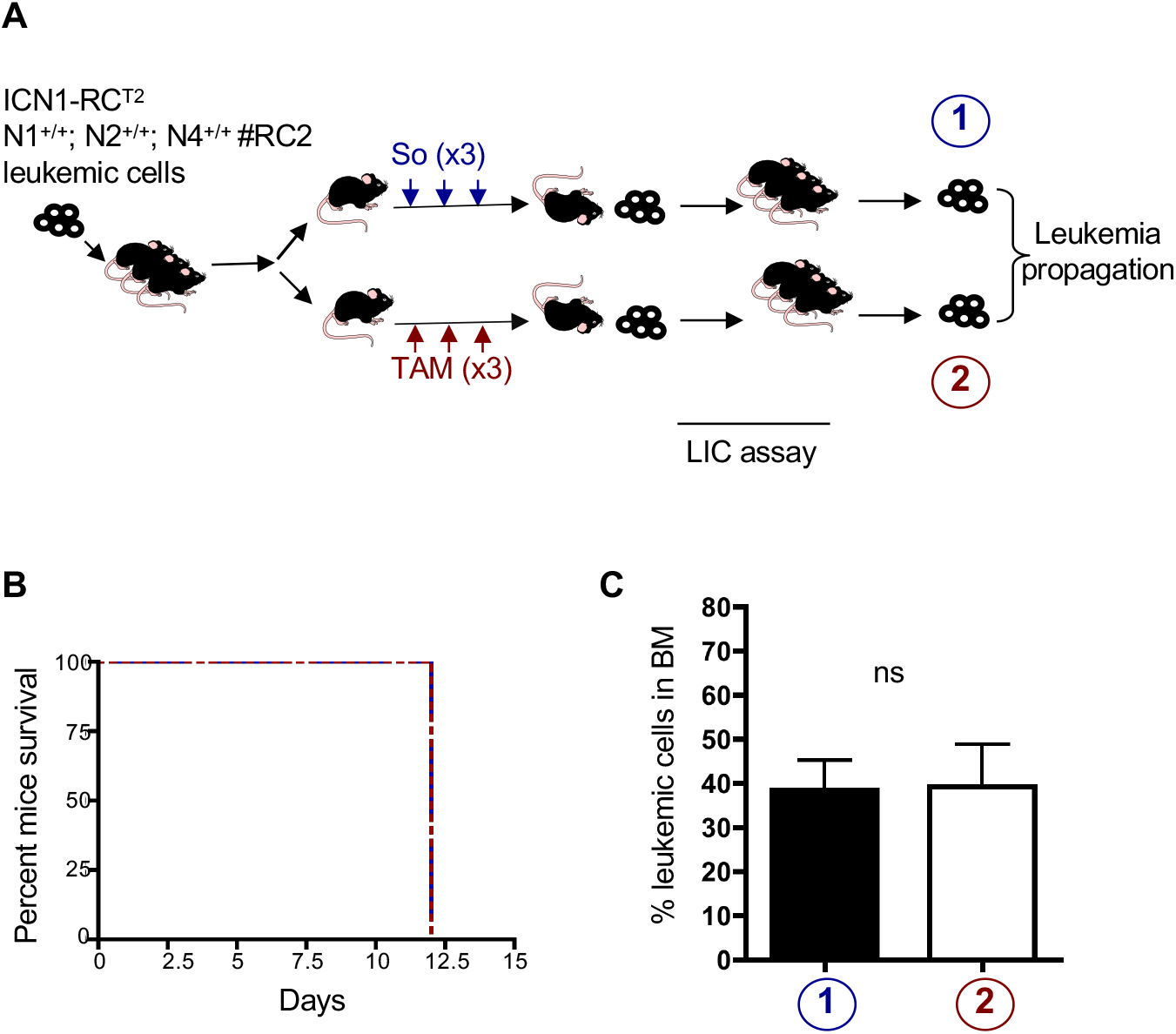
Rosa-Cre activation does not affect leukemia propagation after transplantation. **(A)** Schematic representation of the experiment: Mice were injected with leukemic cells obtained from T-ALL ICN1; RC^T2^; NFAT1^+/+^; NFAT2^+/+^; NFAT4^+/+^. When BM leukemia burden reached about 10-15% leukemic cells in recipients, mice received 3 successive daily injection of either carrier solvent (So, n=3) or tamoxifen (TAM, n=6). Terminally ill mice from both groups were sacrificed 2 days later and leukemic cells from So-treated (blue label, 1) or Tam-treated (red label, 2) cells (10^6^ cells/mouse) were transplanted in wild-type secondary recipients that were followed for leukemia recurrence. **(B)** Kaplan-Meier survival curve of recipient mice infused with 1×10^6^ T-ALL #RC2 cells. Mice were followed overtime for tumor recurrence and recipient mice survival. **(C)** Leukemic burden analysis of recipient mice (n=3 for each group) infused with T-ALL #RC2 cells expressing NFAT factors *Nfat*-proficient (data are represented as ± SEM; n=3; Student’s t test; ns: non significant).

**Supplementary Figure 3.**
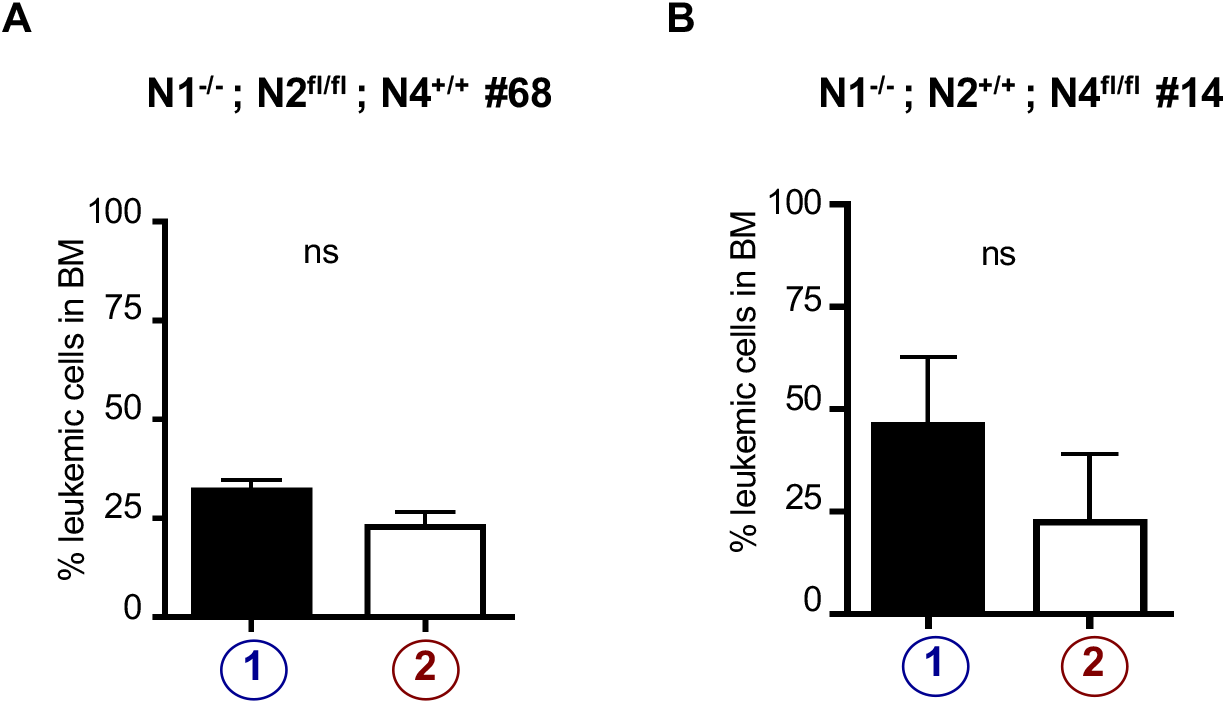
Leukemic load in mice injected with ICN1-induced T-ALL driven only by NFAT2 or NFAT4. T-ALL #68 and #14 expressing only NFAT4 or NFAT2 respectively were generated as described in Fig 4A. Mice were sacrificed when terminally ill, 2 days after the end of So and TAM treatments (1 and 2 in Fig. 5A) and analyzed for leukemic burden (% GFP+ cells for tumor #68 in panel **A**; % tNGFR+ cells for tumor #14 in panel **B).**

**Supplementary Figure 4.**
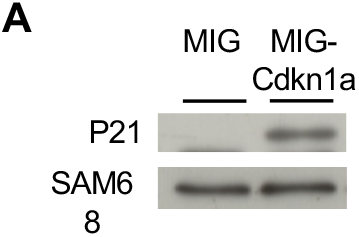
Leukemic cells from T-ALL #3 were transduced with MIG vectors encoding CDKN1A or the control MIG vector without insert. Leukemic cells co-cultured on MS5 stromal cells for 2 days were analyzed by western blot for P21 expression. SAM68 is used as loading control.

**Supplementary Table 1.**

| Leukemia id | Treatment of donor mice | Number of cells injected in recipients | Number of injected recipients | Number of leukemic recipients (time to death, days) | Statistics | % leukemic cells in the BM ± SEM of recipients |
| --- | --- | --- | --- | --- | --- | --- |
| #RC2 | So | 4.10<sup>6</sup> | 3 | 3 (14 ;16 ;16) | ns | 57,3±13,3 |
|  | TAM | 4.10<sup>6</sup> | 3 | 3 (16 ;16 ;18) |  | 56,2±12 |
|  | So | 1.10<sup>4</sup> | 3 | 3 (21 ;21 ;24) | ns | 74,8±2,2 |
|  | TAM | 1.10<sup>4</sup> | 3 | 3 (21 ;23 ;23) |  | 69,78±9,9 |
|  | So | 1.10<sup>3</sup> | 3 | 3 (25 ;25 ;25) | ns | 64,8±13,3 |
|  | TAM | 1.10<sup>3</sup> | 3 | 3 (25 ;28 ;28) |  | 79,5±1,8 |
|  | So | 1.10<sup>2</sup> | 3 | 3 (30 ;32 ;36) | ns | 79,3±0,8 |
|  | TAM | 1.10<sup>2</sup> | 3 | 3 (30 ;34 ;36) |  | 75,6±1,8 |

**Supplementary Table 2.**
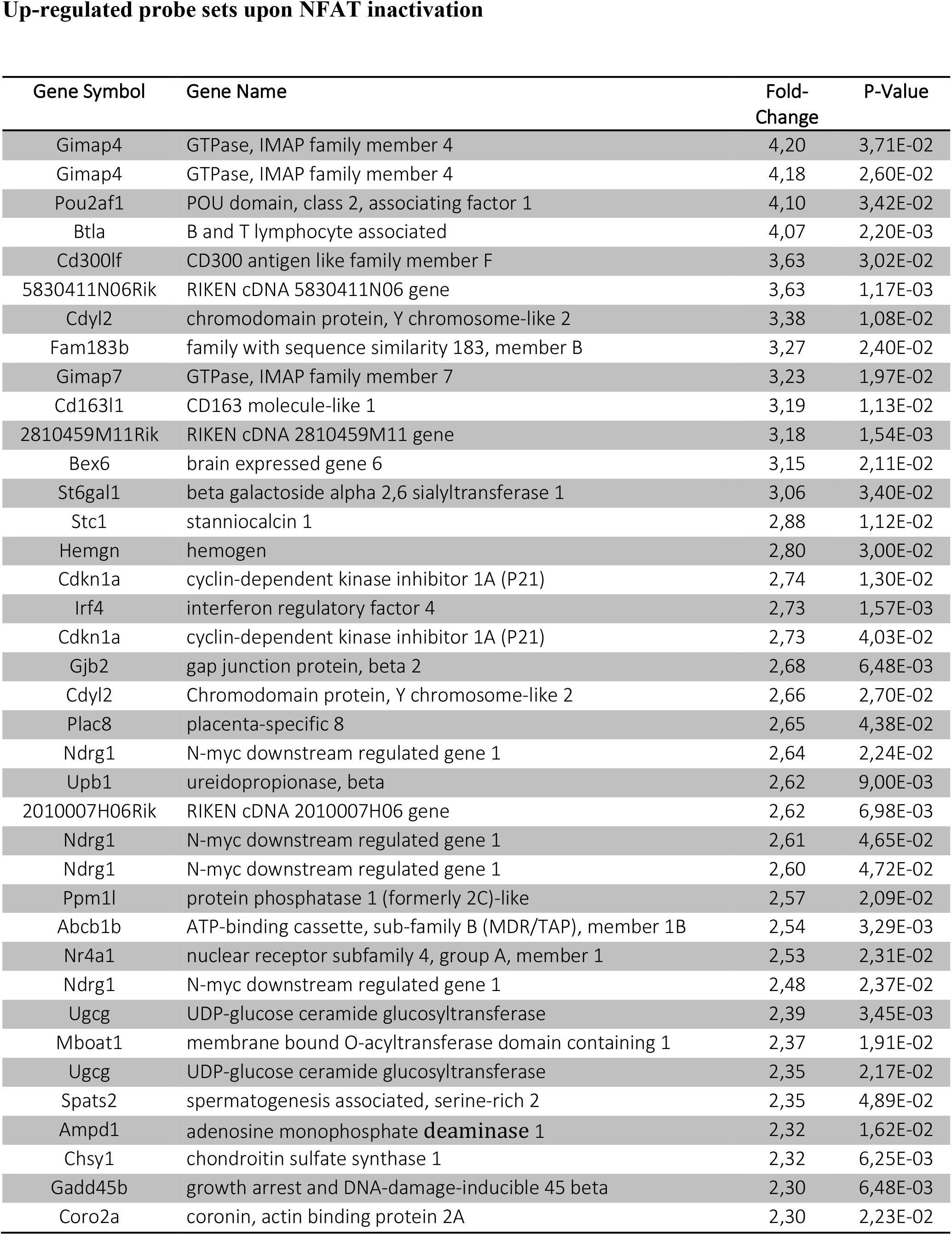
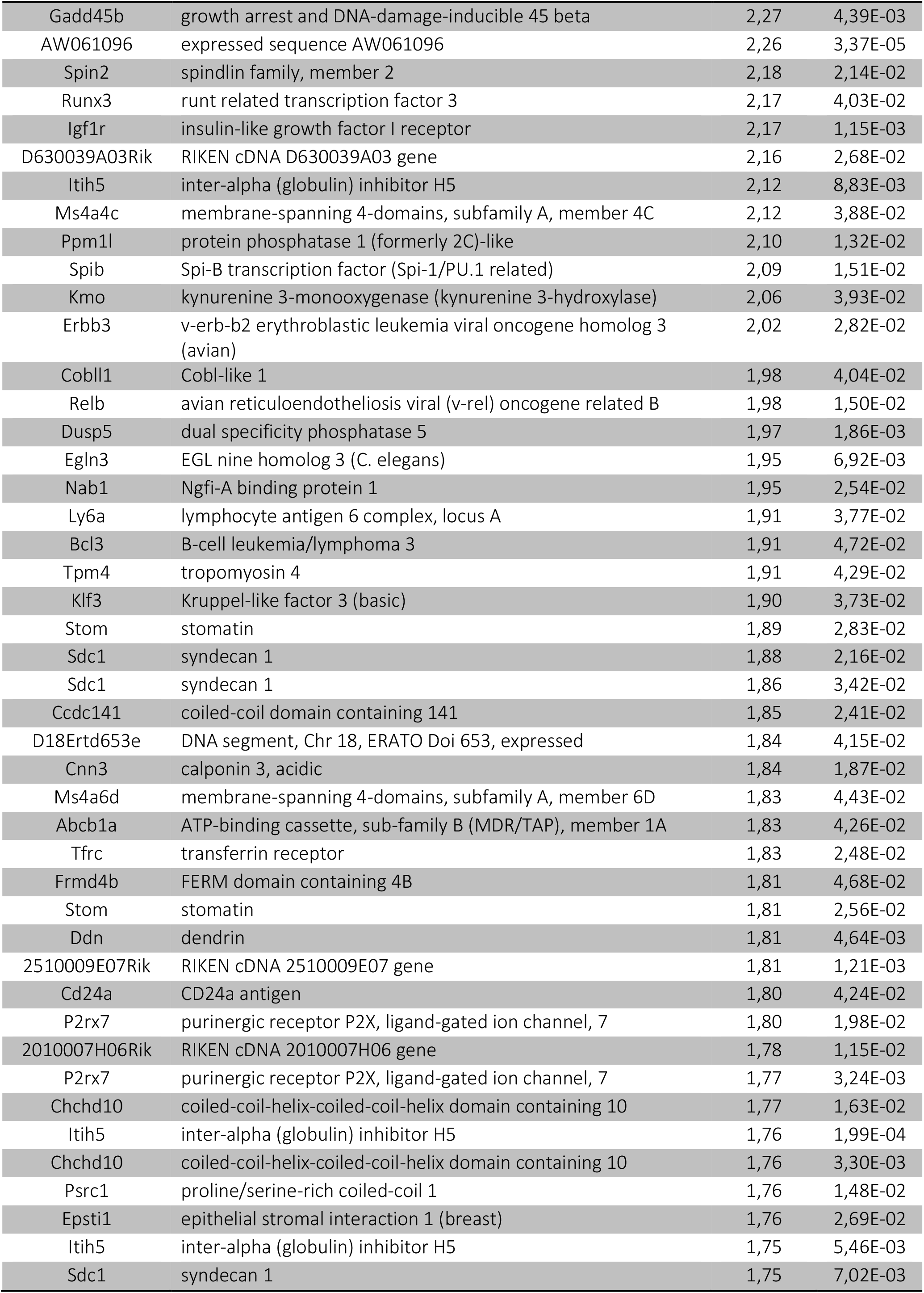
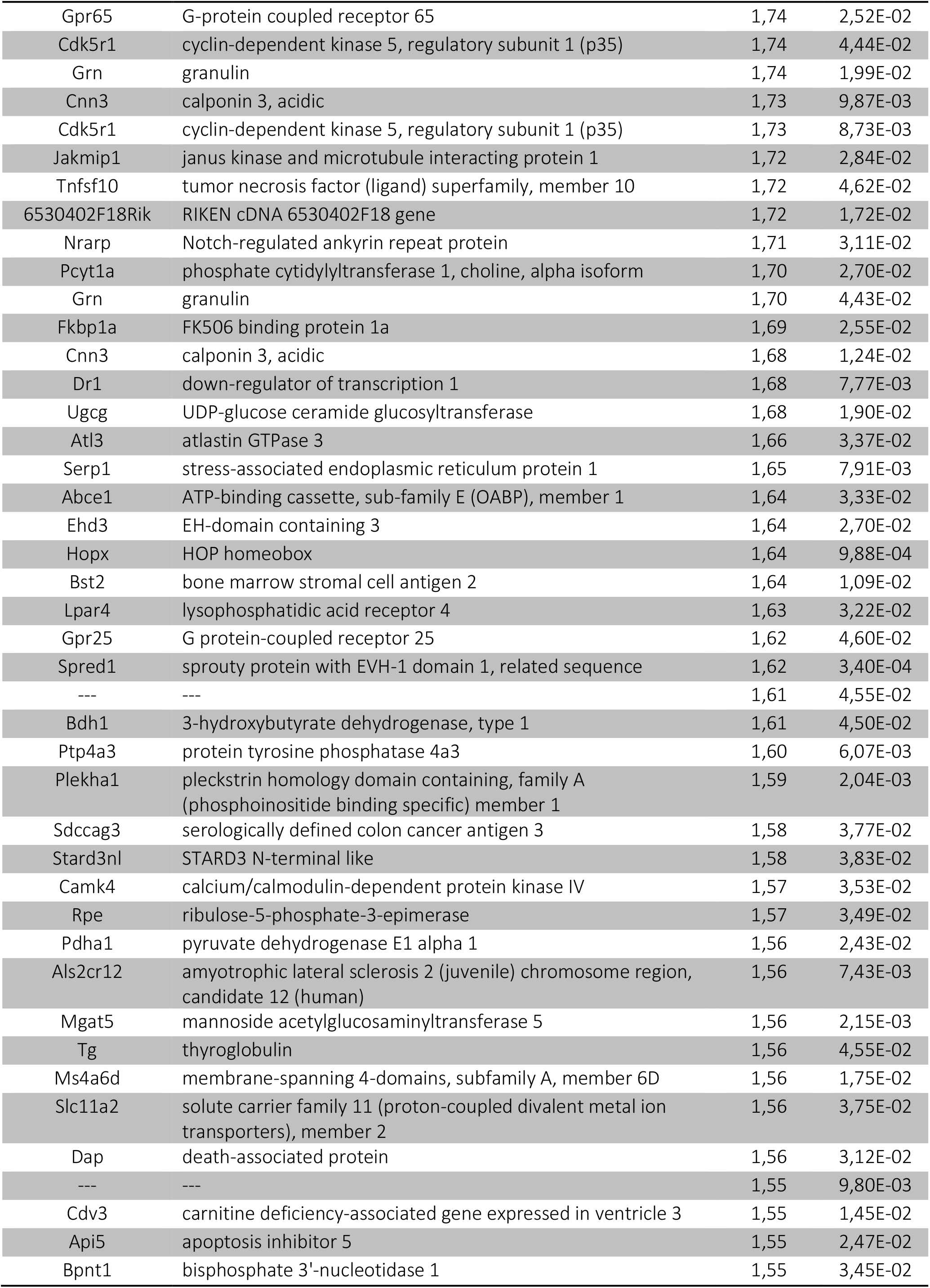
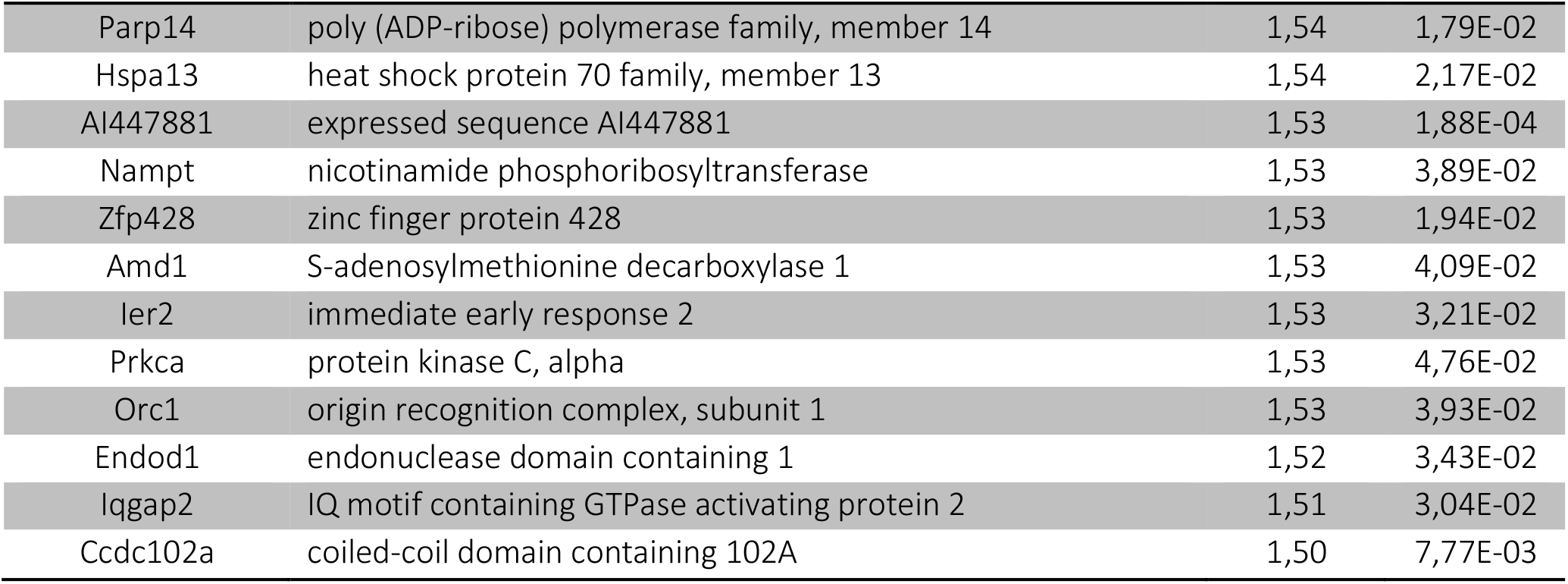
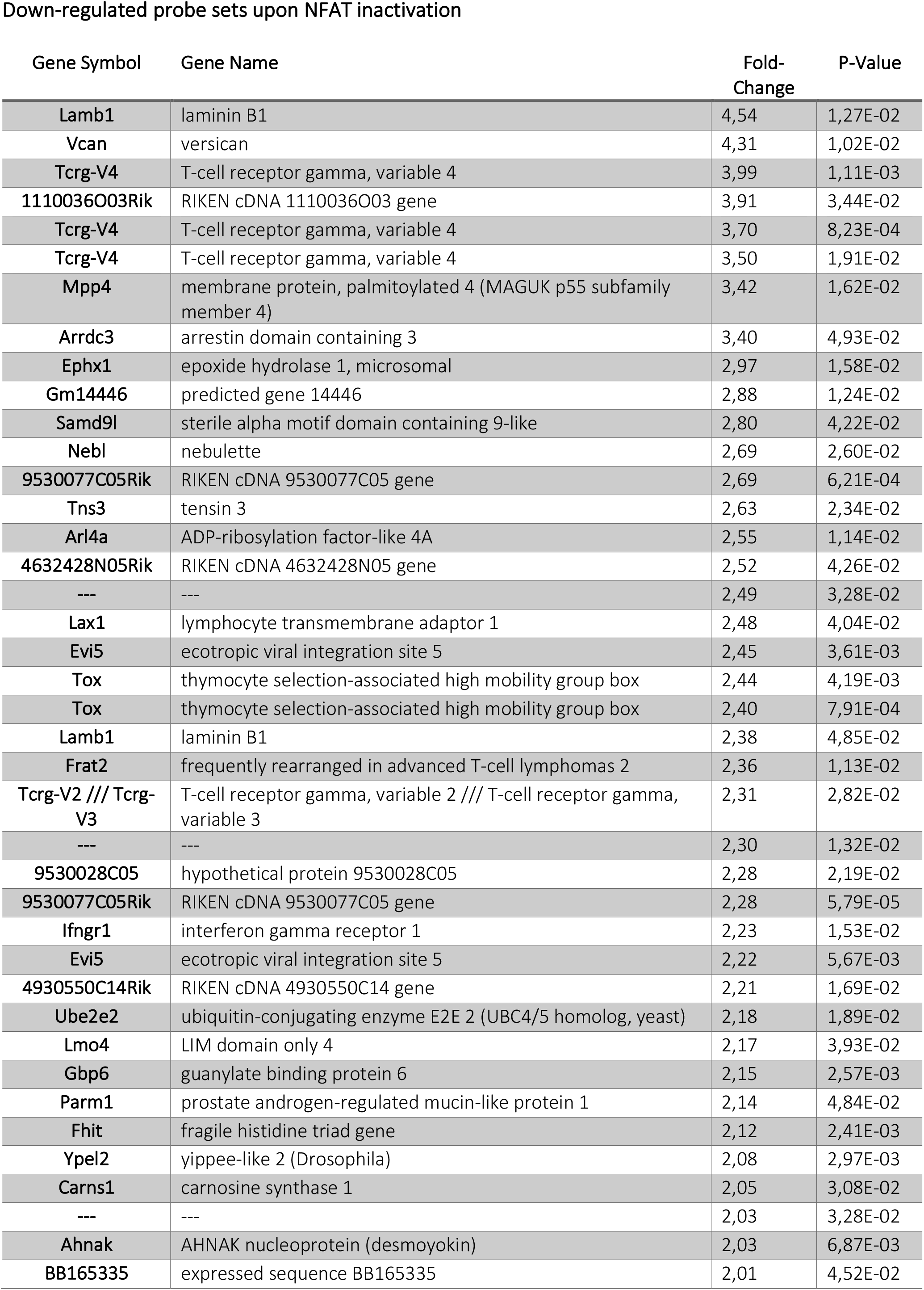
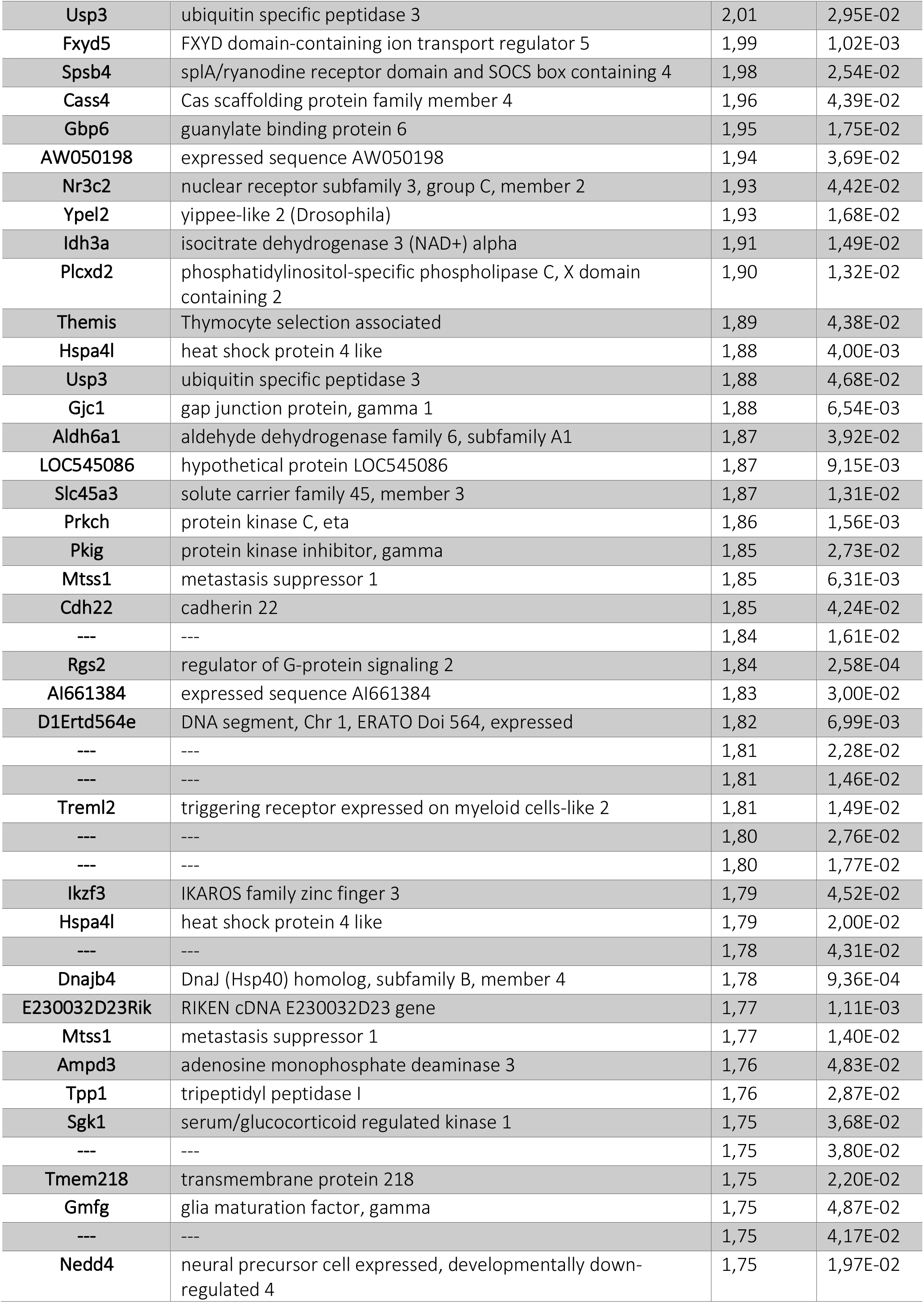
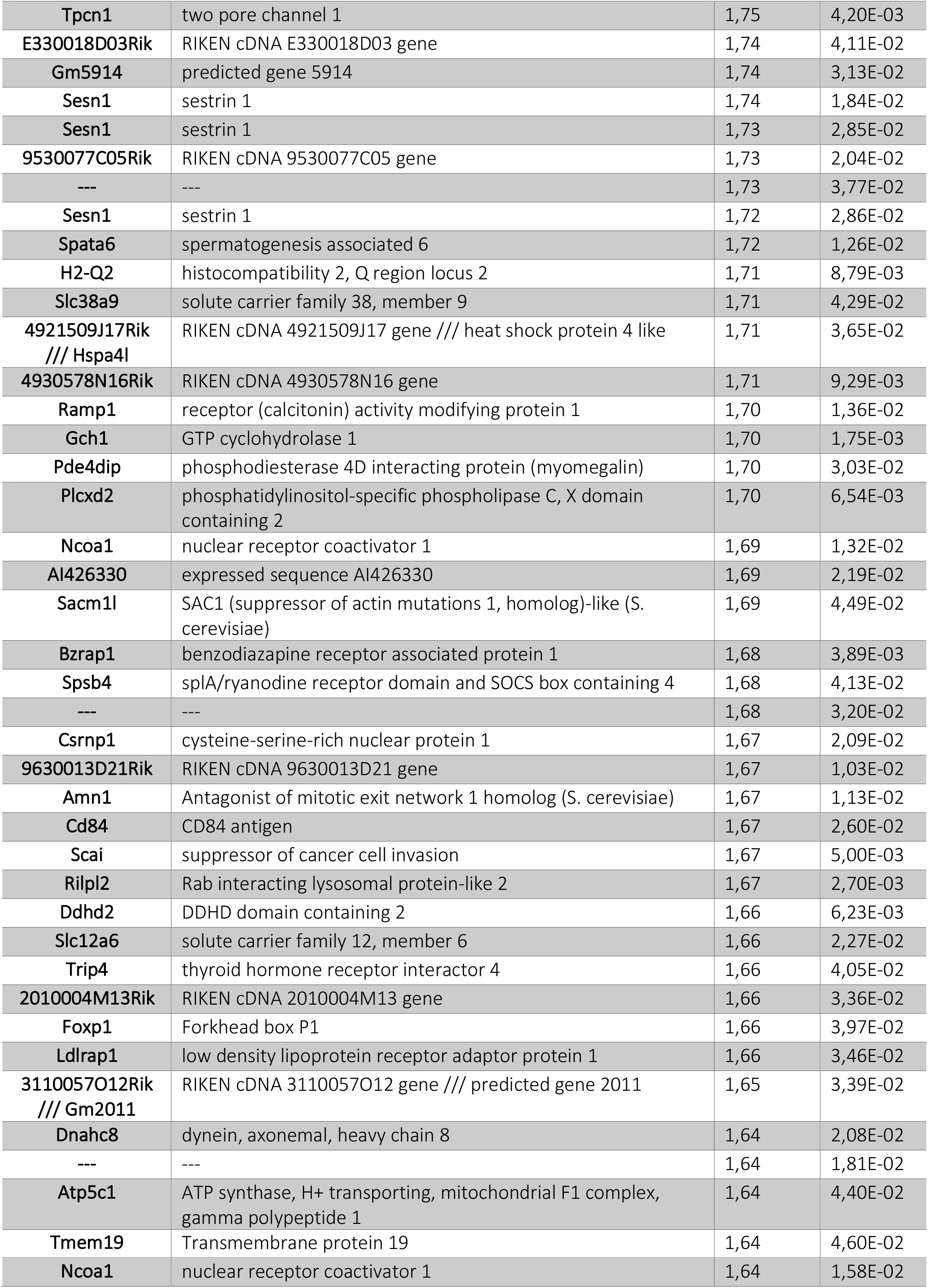
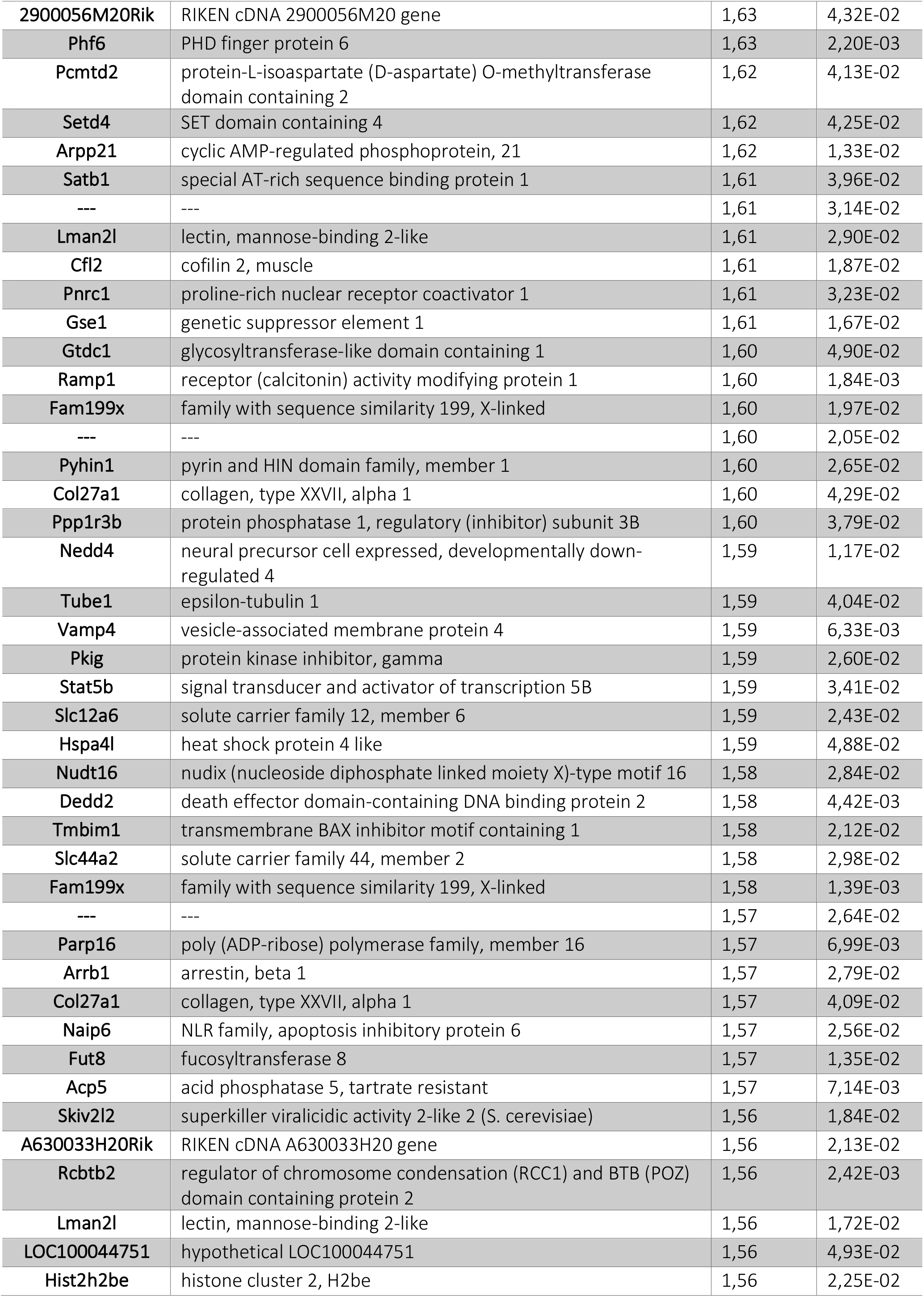
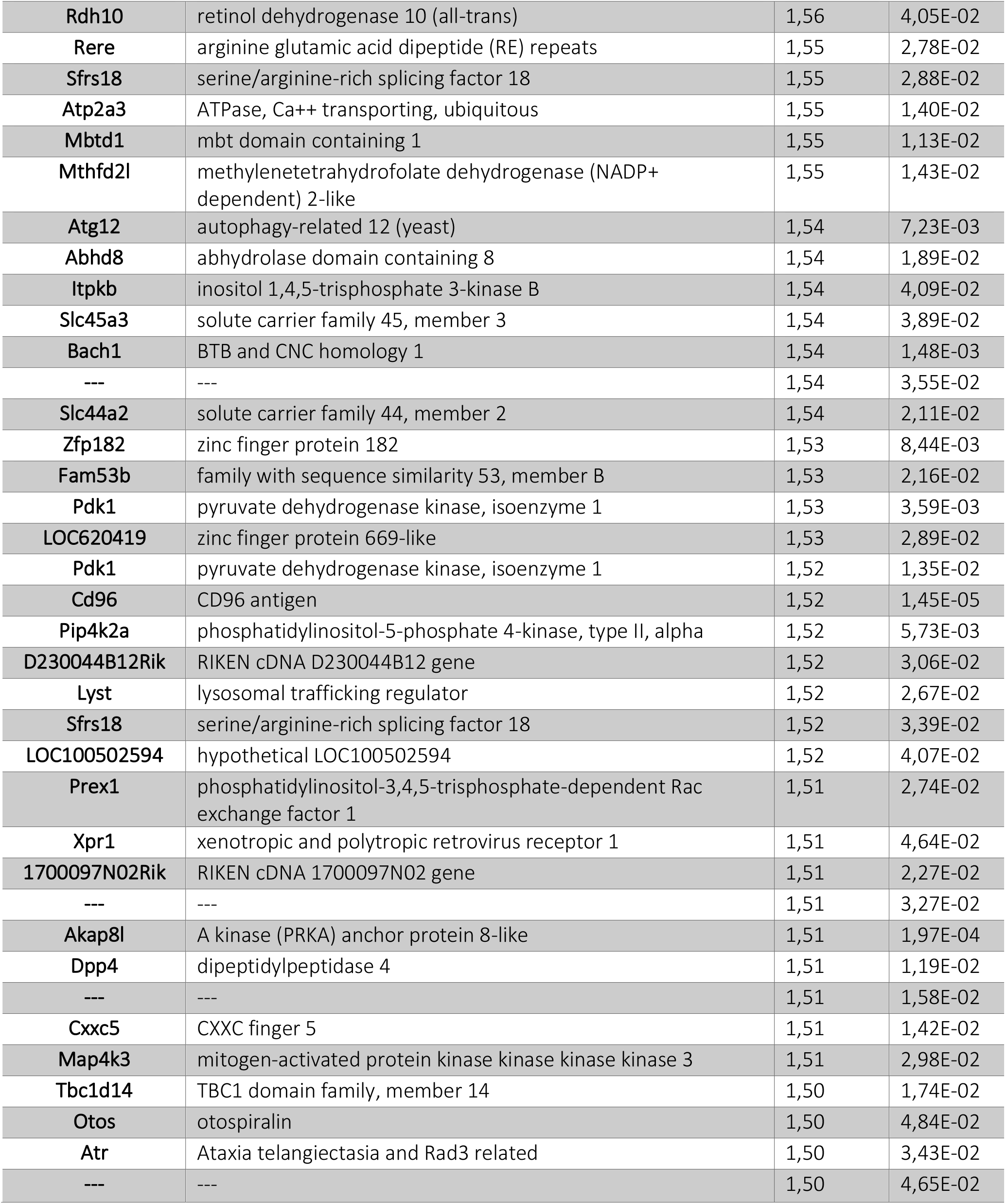
NFAT-dependent transcriptome in T-ALL.

**Supplementary Table 3.**
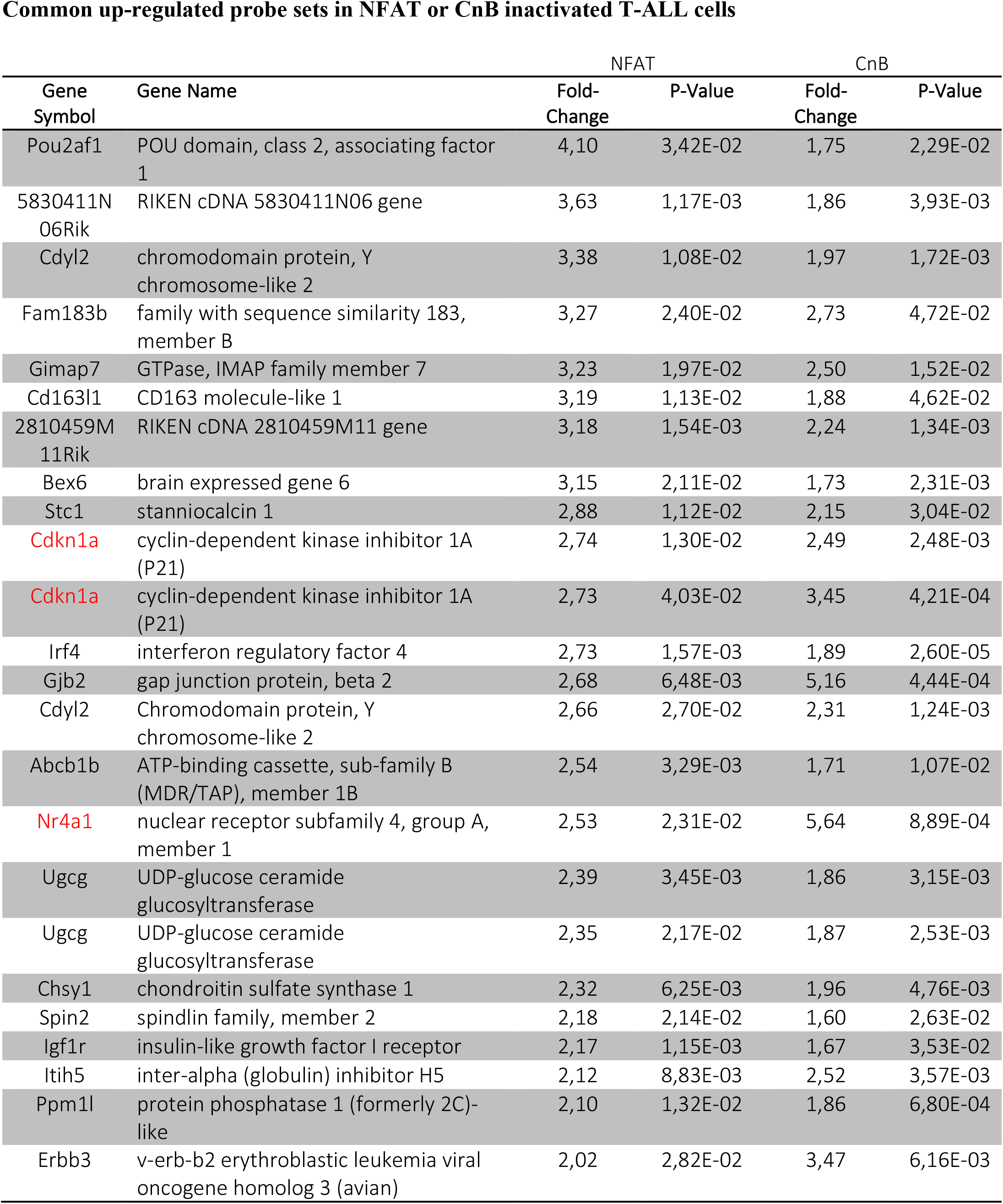
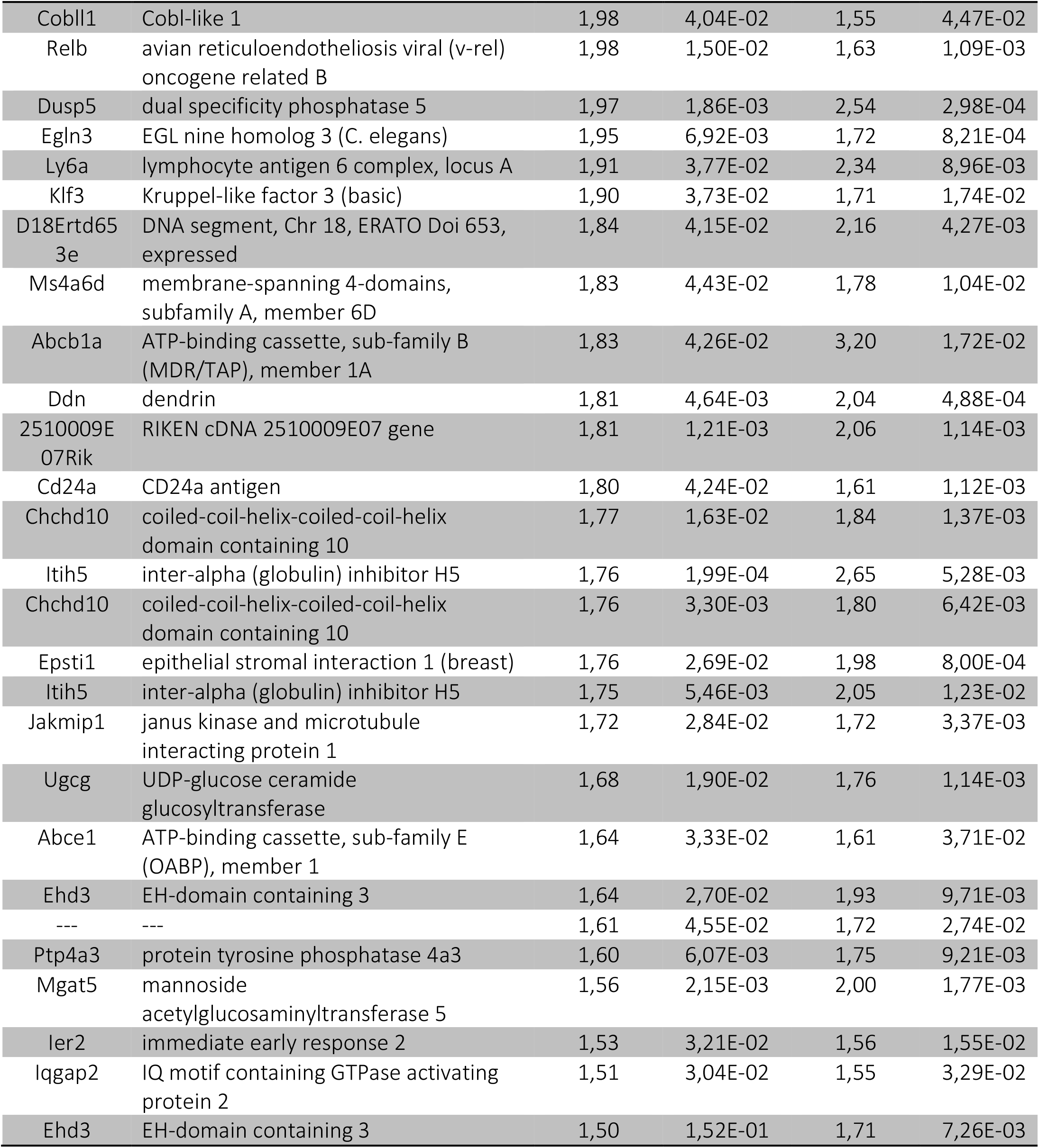
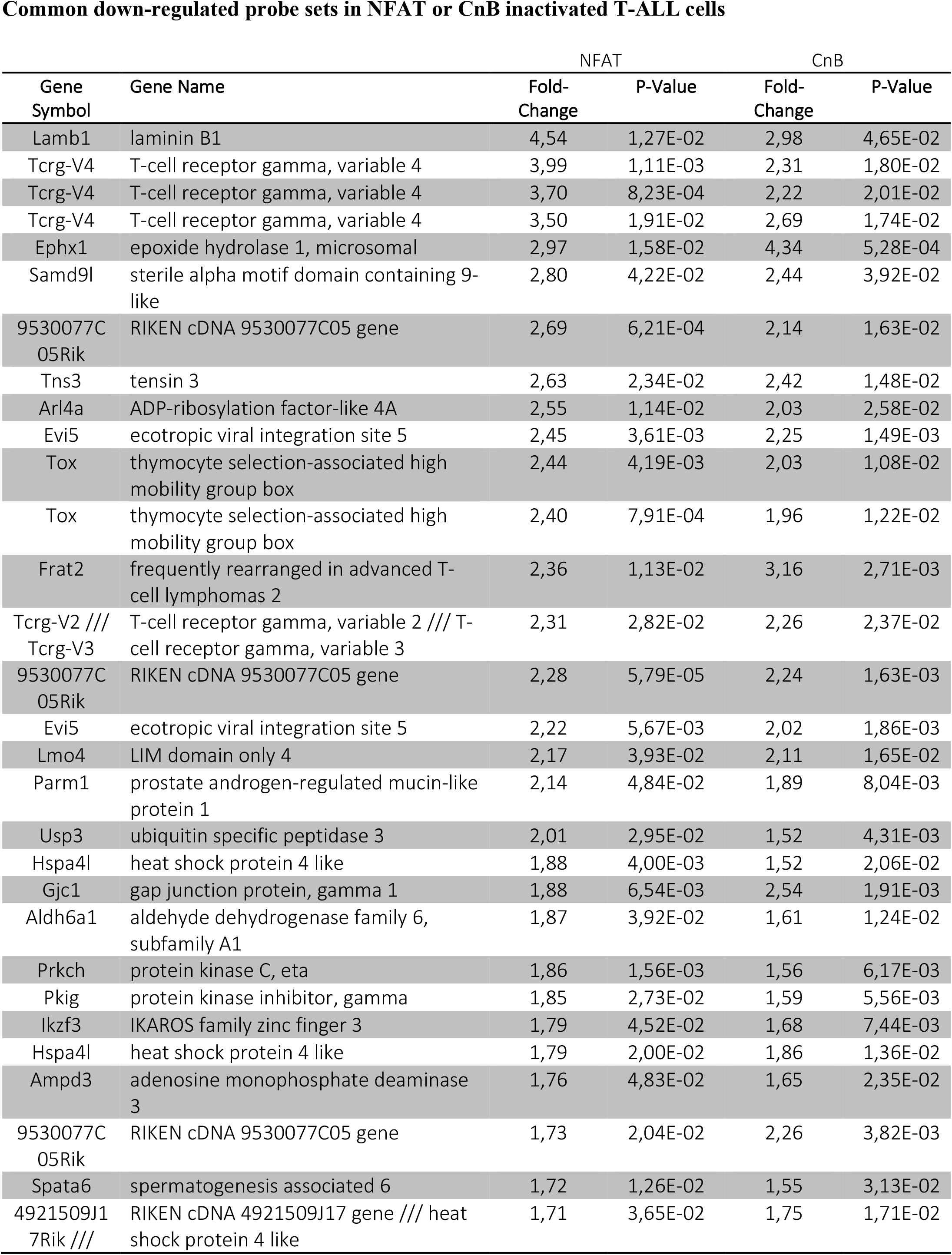
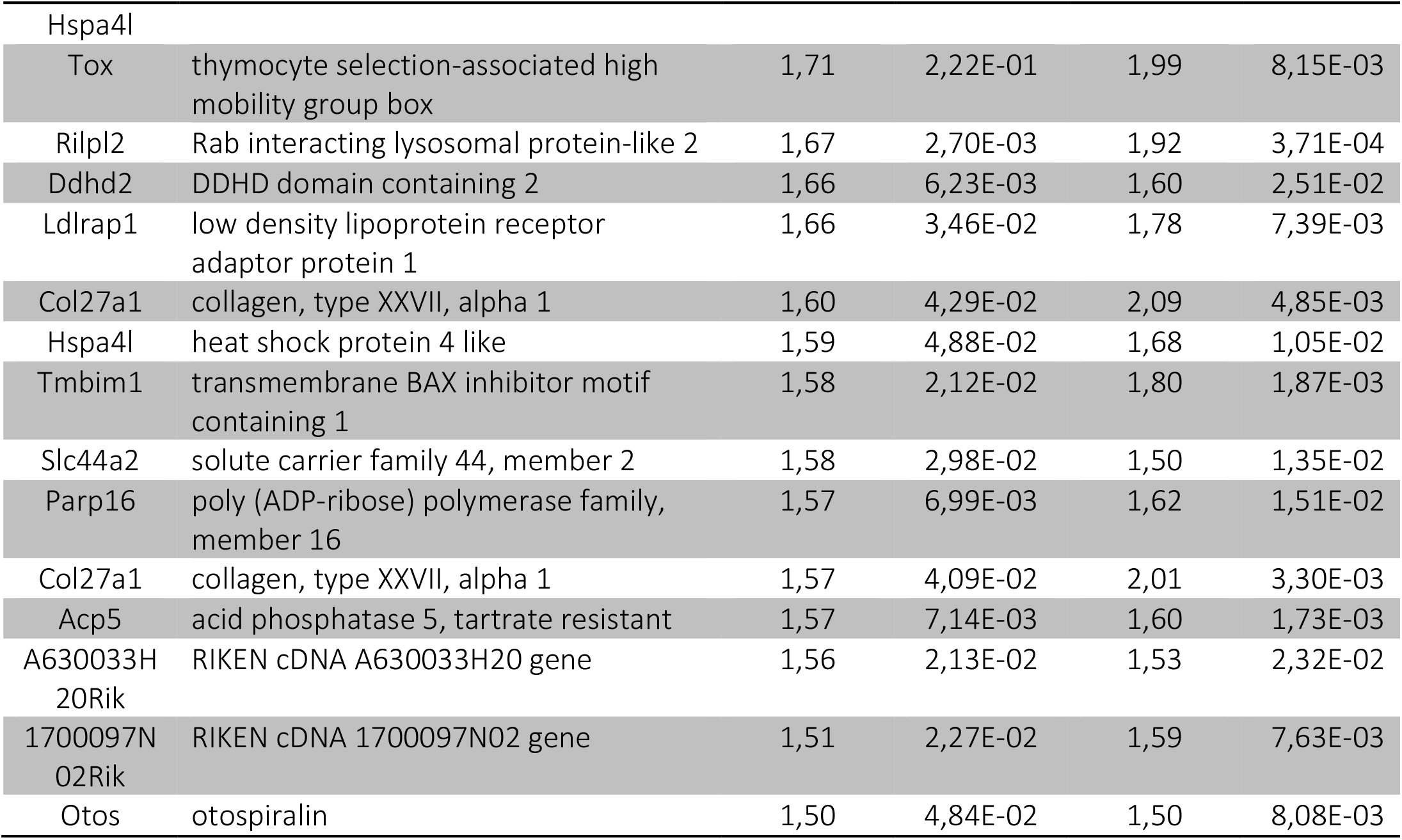
Overlap between NFAT- and Calcineurin-dependent transcriptomes.

## SUPPLEMENTAL MATERIELS AND METHODS

### Cell culture

MS5 (mouse) bone marrow-derived stromal cells were maintained in aMEM (Invitrogen) supplemented with 10% heat-inactivated fetal bovine serum (FBS) (Invitrogen). NIH-3T3 (mouse) cells were maintained in DMEM (Invitrogen) supplemented with 10% heat-inactivated donor bovine serum (Invitrogen). 293T and PlatE cells were maintained in DMEM (Invitrogen) supplemented with 10% FBS (Invitrogen). Leukemic cells were co-cultivated in RPMI (Invitrogen) supplemented with 15% FBS, IL2 and IL7 (10ng/ml; Peprotech). *In vitro* activation of RC^T2^ was performed by a 24h pulse treatment with 4-hydroxytamoxifen (4OHT, 1μM, Sigma Aldrich).

### Retroviral-mediated gene transfer

The cDNAs encoding *CnB1 (PPP3R1)*^8^, and the constitutively active mutant NFAT1 [2+5+8] and NFAT1 [2+5+8]-NLS (a generous gift of Dr A Rao)^21^, were subcloned into MigR1, allowing their co-expression with GFP. Retroviral stocks were obtained by transfection of the PlatE packaging cell line with the desired retroviral vector using Lipofectamine, following manufacturer’s instructions (Lipofectamine 2000, Invitrogen). After 48 hours, the retroviral supernatant was collected and titrated on NIH3T3 cells. tNGFR^+^ ICN1 leukemic cells were spin-infected (1800g for 2 h at 30°C) at the same multiplicity of infection in the presence of 4μg/ml polybrene (Sigma-Aldrich). Following infection, leukemic cells were co-cultured on MS5 for 48-72 hours before flow cytometry sorting or immediately infused into syngeneic mice (1×10^6^ cells/mouse) and expanded *in vivo* for further studies

### PCR genotyping, RNA extraction and RT-PCR analyses

Detection of the different alleles of Nfat1, *Nfat2* and *Nfat4* in genomic DNA of compound mice and leukemic cells was by PCR, using the following primers:

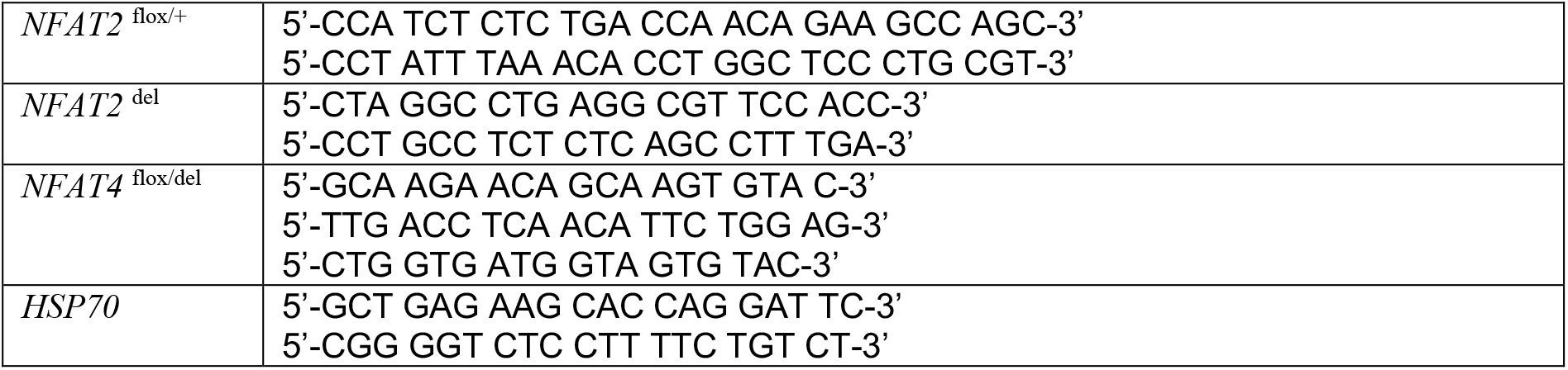

Total RNA was extracted using the RNeasy kit (QIAGEN) and reverse transcribed using random primers and the kit ImProm II Reverse Transcription System (Promega) according to the manufacturer’s instructions. The following primers were used for RT-PCR:

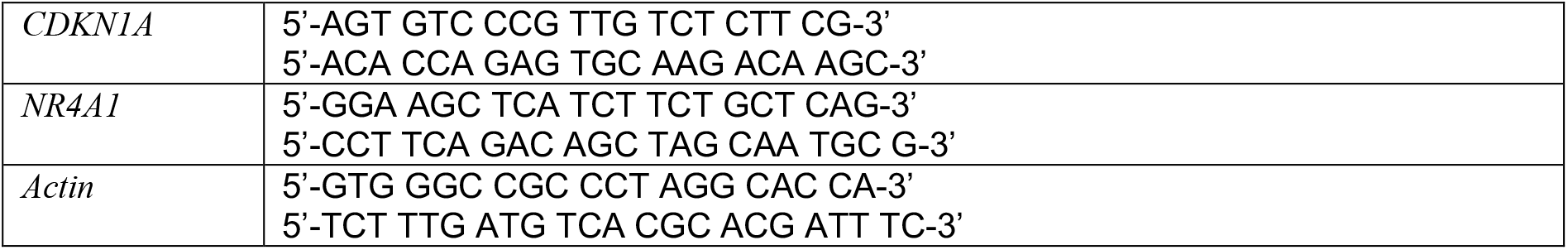

### Western blot

Western blot analyses were performed as described^6^, using antibodies to NFAT2 (sc-7294), NFAT1 (sc-7296), NFAT4 (sc-8321), STAT5 (sc-835) and SAM-68 (sc-333), all from Santa Cruz Biotechnology.

